# Synaptic vesicles transiently dock to refill release sites

**DOI:** 10.1101/509216

**Authors:** Grant F Kusick, Morven Chin, Sumana Raychaudhuri, Kristina Lippmann, Kadidia P Adula, Edward J Hujber, Thien Vu, M Wayne Davis, Erik M Jorgensen, Shigeki Watanabe

**Affiliations:** Department of Cell Biology, Johns Hopkins University, School of Medicine, Baltimore, MD 21205, USA; Biochemistry, Cellular and Molecular Biology Graduate Program, Johns Hopkins University, School of Medicine, Baltimore, MD 21205, USA.; Neurobiology Course, The Marine Biological Laboratory, Woods Hole, MA 02543, USA; Department of Biology and Howard Hughes Medical Institute, University of Utah, Salt Lake City, UT 84112-0840, USA; Solomon H. Snyder Department of Neuroscience, Johns Hopkins University, School of Medicine, Baltimore, MD 21205, USA

**Keywords:** Synaptic vesicle exocytosis, neurotransmitter release, multivesicular release, asynchronous release, synaptic vesicle docking, transient docking, flash-and-freeze, zap-and-freeze

## Abstract

Synaptic vesicles fuse with the plasma membrane to release neurotransmitter following an action potential, after which new vesicles must ‘dock’ to refill vacated release sites. To capture synaptic vesicle exocytosis at cultured mouse hippocampal synapses, we induced single action potentials by electrical field stimulation then subjected neurons to high-pressure freezing to examine their morphology by electron microscopy. During synchronous release, multiple vesicles can fuse at a single active zone; this multivesicular release is augmented by increasing extracellular calcium. Fusions during synchronous release are distributed throughout the active zone, whereas fusions during asynchronous release are biased toward the center of the active zone. Immediately after stimulation, the total number of docked vesicles across all synapses decreases by ∼40%. Between 8 and 14 ms, new vesicles are recruited to the plasma membrane and fully replenish the docked pool in a calcium-dependent manner, but docking of these vesicles is transient and they either undock or fuse within 100 ms. These results demonstrate that recruitment of synaptic vesicles to release sites is rapid and reversible.

---

Synaptic vesicle fusion takes place at a specialized membrane domain: the active zone^1^. The active zone is organized into one or more release sites, individual units at which a single synaptic vesicle can fuse^2^. Ultrastructural studies demonstrate that some synaptic vesicles are in contact with the plasma membrane in the active zone and define the ‘docked’ pool^3, 4^. Since both docking and physiological readiness require engaged SNARE proteins^4–6^, docked vesicles are thought to represent fusion-competent vesicles. In fact, previous studies demonstrate that docked vesicles are partially depleted following stimulation^7–10^. However, it is not clear how release sites are refilled by vesicles to sustain neuronal activity.

Docking of vesicles to refill release sites must be rapid. A single action potential consumes some docked vesicles, bursts of action potentials would be expected to deplete all docked vesicles. Nevertheless, some central synapses can fire at a frequency of one kilohertz^11, 12^. Studies using electrophysiology and electron microscopy indicate that recovery of the docked and readily-releasable vesicle pools is slow –about 3 seconds^7, 8, 13^. However, an emerging body of work suggests that vesicle replenishment constitutes several kinetically and molecularly distinct steps, some of which may occur on very fast timescales^14^. In two notable recent examples, modeling based on physiological data predicted that vesicles reversibly transition from “replacement sites” to “docking sites” within milliseconds of an action potential^15, 16^, and experiments with flash-and-freeze electron microscopy demonstrated that Synaptotagmin-1 mutants with docking defects can be reversed by binding calcium^9^. These fast vesicle docking events have been proposed to correspond to calcium-induced changes between loose and tight assembly of the SNARE complex, which may be both very fast and reversible^17^. However, there is currently no ultrastructural evidence for such fast and reversible docking steps at wild-type synapses.

To characterize the ultrastructure of vesicle docking and fusion at active zones, we developed a method to trigger single action potentials by electrical stimulation followed by high-pressure freezing at defined time points called ‘zap-and-freeze’. Using this approach, we first characterized the spatial and temporal organization of fusion sites following a single action potential. We observed that during synchronous release, multiple vesicles can fuse per action potential within the same active zone, even in physiological extracellular [Ca^2+^]. Fusions during synchronous release occur throughout the active zone, but during asynchronous release are concentrated at the center of the active zone. We then followed the fate of docked vesicles. Unexpectedly, ∼40% of docked vesicles are lost immediately after stimulation, due not only to fusion but also to undocking. These are then fully replaced by newly docked vesicles within 14 ms, perhaps to counteract short-term depression. This transient docking requires residual calcium in the terminals and only lasts for 100 ms or less. This sequence of rapid redocking and subsequent slow undocking may underlie facilitation.

## Results

### Zap-and-freeze captures synaptic vesicle fusion

To capture exocytosis with millisecond precision under physiologically-relevant conditions, we developed a system to electrically stimulate neurons before high-pressure freezing: a small, portable field-stimulation device with a photoelectric control switch (Fig. 1a). This device can be charged, then loaded into a high-pressure freezer and discharged with a flash of light to generate a 1 ms 10 V/cm stimulus before freezing at defined time points (see Methods).

**Figure 1.**
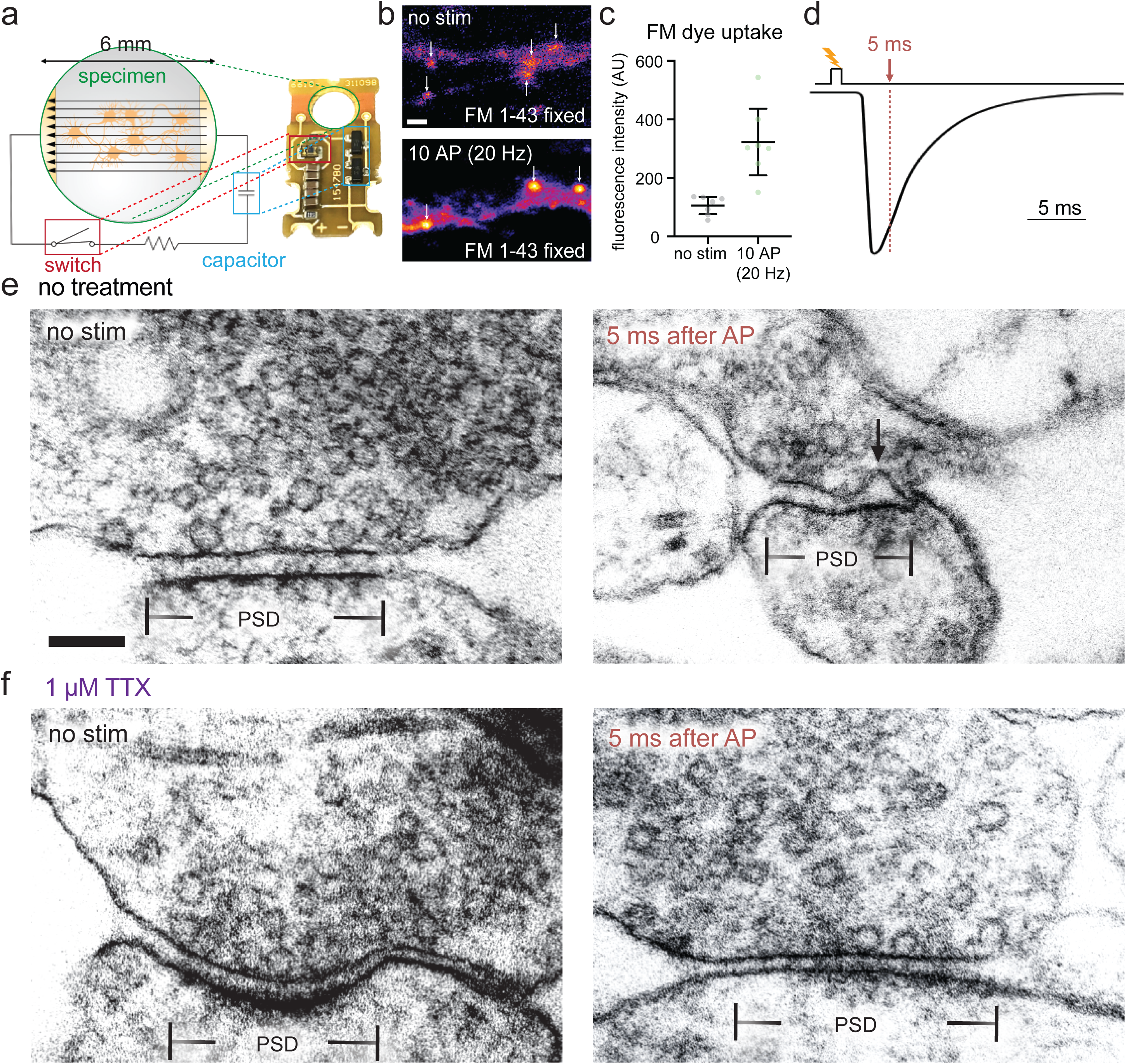
**Zap-and-freeze captures synaptic vesicle fusion. a**, Schematic and photograph of the zap-and-freeze stimulation device. **b**, Epifluorescence micrographs of cultured mouse hippocampal neurons pre-incubated in 30 µM Pitstop 2 in physiological saline (1 mM Ca^2+^) for 2 min, then either not stimulated or subjected to 10x 1 ms pulses at 20 Hz, at 37 °C in FM 1-43FX, followed by washing and fixation. Arrows indicate putative presynaptic terminals, identified by their increased FM labeling relative to the rest of the axon, shape, and size. Scale bar: 2 μm. **c**, Quantification of the experiment described in **b**; n = 7 fields of view with 20 total putative boutons quantified per image; p = 0.003, Welch’s t-test. N = 1 experiment. Note that the images shown in **b** are crops of a small portion of the full field of view for each image. Error bars indicate mean and 95% confidence interval. **d**, Experimental design for stimulation and freezing, showing a diagrammatic excitatory postsynaptic potential for reference (based on ^39^). A 1-ms square pulse is applied to trigger a single action potential, then neurons are frozen 5 ms after the beginning of the pulse (this is the earliest possible freezing time on the high-pressure freezer, see Methods). **e-f**, Transmission electron micrographs of synapses from neurons high-pressure frozen in 1.2 mM Ca^2+^ either **e** without or **f** with tetrodotoxin (TTX), which prevents action potential firing. Samples were frozen either with no stimulation (“no stim”) or 5 ms after stimulation, which presumably initiates an action potential (“5 ms after AP”). The arrow indicates a pit in the active zone, which is presumed to be a synaptic vesicle fusing with the plasma membrane. The active zone is defined as the presynaptic plasma membrane opposite the post-synaptic density (PSD). Scale bar: 100 nm. Electron micrographs are from experiments described in Figure 3. See Supplementary Data Table 1 for full pairwise comparisons and summary statistics.

To test whether this device is functional, we performed FM 1-43 loading experiments in mouse hippocampal neurons cultured on 6-mm sapphire disks. The lipophilic FM dye is taken up by compensatory endocytosis after synaptic vesicle fusion^18^. To prevent destaining by exocytosis, we applied Pitstop 2 (30 µM) 2 minutes prior to loading. Pitstop 2 is a nonspecific^19^ but nonetheless potent inhibitor of clathrin-mediated vesicle formation^20^. The dye is taken up during ultrafast endocytosis, which is clathrin-independent, but will be trapped in synaptic endosomes which are resolved by clathrin-mediated budding^21^. Neurons were stimulated 10 times at 20 Hz, each pulse lasting 1 ms, which induces a single action potential. Following stimulation and fixation, presynaptic terminals were strongly labeled with FM 1-43 (Fig. 1b-c; 3-fold increase relative to no-stim control, p = 0.003, see Supplementary Figure 1c for full fields of view from micrographs), suggesting that the stimulation device triggers action potentials and synaptic activity.

With the stimulation device validated, we next tested whether exocytic intermediates can be captured by high-pressure freezing. Experiments were performed at 37 °C and 1.2 mM external calcium, roughly the [Ca^2+^] of the interstitial fluid in the brain^22^. We applied a single 1 ms pulse, which likely triggers a single action potential^23^. Cells were frozen 5 ms after stimulation (Fig. 1e-f), which is the earliest possible time point given the mechanics of the high-pressure freezer (see Methods). Samples were then prepared for electron microscopy, and images were acquired and quantified blind (see Methods). We defined the active zone as the membrane domain directly apposed to the postsynaptic density (Fig. 1e-f). Typically, the length of the active zone visible in single sections was ∼300 nm, with a wide range including much larger active zones (Supplementary Fig. 4f). We quantified any active zone membrane deflections greater than 10 nm by visual inspection as pits. If similar deflections are found outside the active zone, they are measured but considered endocytic^7^ or membrane ruffles and thereby not included in the data (see Supplementary Figure 1a-b for examples of features quantified as pits or not). We also counted the number of vesicles that were 0 to 100 nm above the plasma membrane within the area of the active zone and classified those that appeared to be in physical contact with the plasma membrane as docked (0 nm from the plasma membrane). In stimulated samples 18% of the synaptic profiles exhibited exocytic pits in the active zone (57/ 316), whereas in unstimulated cells only 2% of the synaptic profiles exhibited pits (6/ 275), and in cells in which action potentials were blocked by tetrodotoxin only 1% of the profiles contained pits (2/256) (Fig. 1f). Thus, the device induces *bona fide* action potentials and vesicle fusion, which can be reliably captured in electron micrographs. By analogy to the previously-developed flash-and-freeze^8^, this technique is called ‘zap-and-freeze’.

### Multivesicular release is prominent in cultured hippocampal neurons

It has long been debated whether univesicular or multivesicular release predominates^24^. From single synaptic profiles, 2% (6/316 synaptic profiles) exhibited multiple pits. Although rare, the presence of multiple pits in the same image indicates that more than one vesicle in an active zone can fuse after a single action potential, an event known as multivesicular release^24^. However, the frequency of such events cannot be determined from single sections, but rather requires reconstruction of whole active zones from serial sections (Fig. 2). To quantify synaptic vesicle fusions per synapse, cultured hippocampal neurons were stimulated in 1.2 mM Ca^2+^ at 37 °C and frozen 5 ms after stimulation. Over 60 active zones were reconstructed for each condition and morphometry performed blind (Supplementary Fig. 2a for example micrographs). In unstimulated samples, 3% of the synapses contained a pit (2/62), whereas in stimulated samples 35% of the synapses exhibited at least one pit (24/68, Fig. 2a). Of those with at least 1 pit, 38% of active zones (9/24) contained multiple pits. All pits ranged from the size of a synaptic vesicle to expected sizes of vesicles at late stages of collapse into the plasma membrane (Fig. 2d; full range of pit widths at base, 24-89 nm). These results suggest that multivesicular release is prominent in cultured hippocampal neurons.

**Figure 2.**
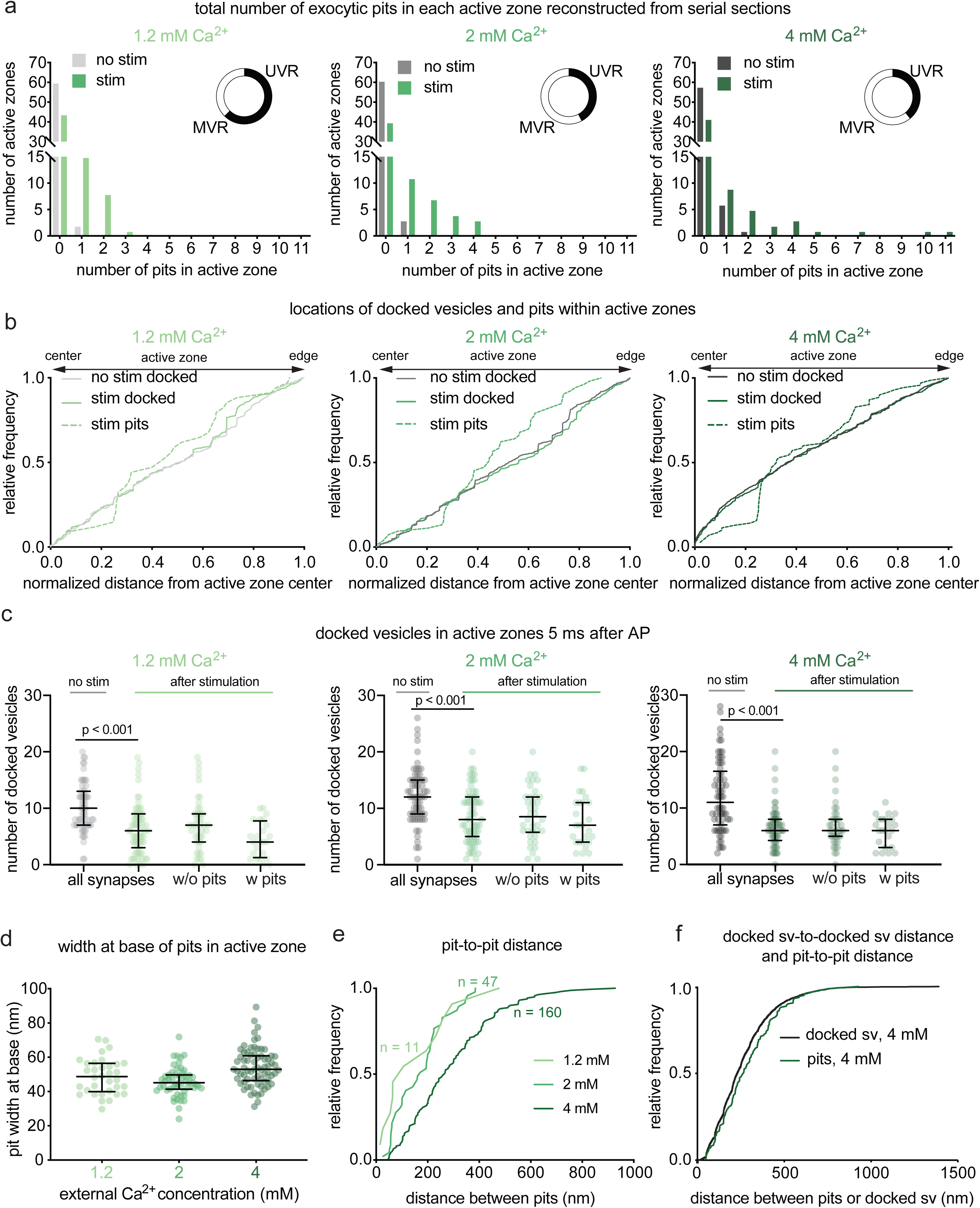
**Multiple fusion events at single active zones after a single action potential. a**, Frequency distributions of number of fusion events 5 ms after an action potential (green) or without stimulation (grey) in solutions of 1.2, 2, or 4 mM Ca^2+^ (1.2 mM, no stim, n = 62; 1.2 mM, stim, n = 68; 2 mM, no stim, n = 64; 2 mM, stim, n = 66; 4 mM, no stim, n = 65; 4 mM, stim, n = 64 reconstructed active zones). Insets show the proportion, out of the active zones that contained at least 1 fusion event, that contained 1 fusion event (UVR, univesicular release) or more than 1 fusion event (MVR, multivesicular release). Fusion events are defined as “pits” in the active zone. Including all active zones from stimulated samples, number of pits was not significantly different in different Ca^2+^ concentrations (p = 0.88); including only synapses with at least 1 fusion event, the numer of pits was significantly greater at 4 mM than at 1.2 mM (p = 0.042). The proportion of synapses that contained at least 1 pit was not different between samples stimulated in different Ca^2+^ concentrations (chi-square = 0.2771, df = 2, p = 0.87). **b,** Cumulative relative frequency distributions of locations of docked vesicles (with and without stimulation) and pits (after stimulation) within the active zone (n = 34, 53, 70 pits; 384, 768, 778 docked vesicles without stimulation; 384, 579, 423 docked vesicles with stimulation, ordered by increasing Ca^2+^ concentration). Locations are normalized to the size of the active zone and to the expected density of objects within a circular area by taking the square of the distance of a pit or vesicle to the center of the active zone divided by the half-length of the active zone: 0.25 would indicate a vesicle or pit halfway between the center and edge. Docked vesicles were not biased toward the center or except for samples frozen in 4 mM Ca^2+^, which were biased toward the center, and samples frozen after stimulation in 2 mM Ca^2+^, which was biased toward the edge (p > 0.1 for each, except for 2 mM Ca^2+^ stim, p =0.02; 4 mM Ca^2+^ no stim and stim, p < 0.001 for each). Vesicles fusions not biased toward the center or the edge (p > 0.9 for 1.2 mM 2 mM, p = 0.05 for 4 mM). For each calcium concentration, the median location of pits and docked vesicles in the active zone after stimulation were similar to those of docked vesicles from no-stim controls (p > 0.9 for each). **c**, number of docked vesicles in each active zone reconstruction 5 ms after an action potential (green) or without stimulation (grey); same n as in **a.** The number of docked vesicles was not significantly different between synapses with and without pits for each calcium concentration (p > 0.1 for each comparison; Kruskal-Wallis test with post-hoc Dunn’s multiple comparisons). Vesicles that appeared to be in contact with the plasma membrane were considered docked. **d**, Width at base of pits in the active zone. Error bars indicate median and interquartile range. **e**, Cumulative relative frequency distributions of distances from center to center of pits within the same active zone, sorted by external calcium concentration. **f**, Cumulative relative frequency distributions of distances from center to center of pits and docked vesicles within the same active zone (n = 218 pairs of pits, 5438 pairs of docked vesicles; see Methods for description of distance calculation). Pit-to-pit distances were slightly greater than docked vesicle-to-docked vesicle distances (pits median: 224 nm, docked median: 265 nm; p = 0.02). Pairs of pits were from stimulated samples in 4 mM Ca^2+^; pairs of docked vesicles were from the no-stim 4 mM Ca^2+^ experiment. Scale bar: 100 nm. PSD: post-synaptic density. AP: action potential. Error bars in **c** indicate median and interquartile range. All data are from two experiments from separate cultures frozen on different days.; experiments in 1.2 mM Ca^2+^ were performed on separate days from a separate culture from the experiments in 2 mM and 4 mM Ca^2+^. Number of pits and docked vesicles per active zone was compared using Kruskal-Wallis tests with post-hoc Dunn’s multiple comparisons test. For pits, full pairwise comparisons were performed; for docked vesicles, only numbers of vesicles before and after stimulation at each Ca^2+^ concentration were compared. Proportions of active zone reconstructions that contain at least one pit were compared using a chi-squared test. Bias of pit locations toward the center or edge of the active zone was tested by comparing each group to a theoretical median of 0.5, the expected median for a random distribution, using two-tailed one-sample Wilcoxon signed-rank tests. Locations of pits, docked vesicles after stimulation, and docked vesicles without stimulation were compared using a Kruskal-Wallis test followed by post-hoc Dunn’s test between pits and no-stim docked vesicles and stim docked vesicles and no-stim docked vesicles for each calcium concentration. P-values from all these pairwise and one-sample comparisons were adjusted with Bonferroni correction accounting for the total number of tests. The distributions of distances between pits in the same active zone and docked vesicles within the same active zone were compared using a two-tailed Wilcoxon rank-sum test. See Supplementary Data Table 1 for full pairwise comparisons and summary statistics. See Supplementary Data Table 2 for summary statistics of docked vesicle and pit counts for each experimental replicate.

### Multivesicular release is augmented by increasing extracellular calcium

To further assess the number of release sites per active zone, we enhanced release probability by increasing the extracellular calcium concentration^25^ from 1.2 mM to 2 mM and 4 mM calcium. Fusion was assessed by the presence of pits in the reconstructed active zones. Increasing the extracellular Ca^2+^ concentration did not change the fraction of synapses that responded (Fig. 2a): at all calcium concentrations only ∼35% of active zones exhibited fusion pits (pits per active zone: 1.2 mM 35% 24/68; 2 mM 39% 26/66; 4 mM 34% 23/64; p = 0.87). However, increasing calcium did augment multivesicular release. In 1.2 mM Ca^2+^ 38% of active zones containing at least one fusion exhibited multivesicular release, in 2 mM Ca^2+^ 58% exhibited multivesicular release, and in 4 mM Ca^2+^ 61% exhibited multivesicular release, including one active zone with 11 pits (Supplementary Fig. 2). The presence of pits in ∼35% of synapses at all calcium concentrations is consistent with the fraction of high-release probability terminals in hippocampal synapses^26–29^, as well as potentially ‘silent’ synapses that never respond to stimulation^30^.

### Release events can be coupled

The presence of several pits within single active zones suggests that each synapse likely has more than one release site. This is consistent with the localization pattern of many proteins essential for neurotransmitter release, including calcium channels. These proteins are clustered, and several clusters seem to be distributed throughout the active zone^31–34^. To assess the distribution of release sites within active zones at the ultrastructural level, we mapped the locations of docked vesicles and exocytic pits (example maps and sizes in Supplementary Fig. 3a-b). At low calcium concentrations, fusing vesicles were often found adjacent to each other (Supplementary Fig. 1, 2), suggesting that neighboring vesicles fuse simultaneously (Fig. 2e). At 1.2 mM Ca^2+^ pits were often within ∼100 nm of each other (median 106 nm, n = 11 pairs).

However, with increasing calcium concentrations, fusion events were less tightly coupled. Adjacent fusions were still observed but additional pits were dispersed across the active zone (Fig. 2e; 2 mM Ca^2+^, median 171 nm, n = 47 pairs; 4 mM Ca^2+^, median 265 nm, n = 160 pairs). At 4 mM Ca^2+^, the median distance between pits was roughly similar to the distance between docked vesicles (Fig. 2f, docked = 229 nm; pits = 265 nm; p = 0.02). Thus, at high calcium concentrations, release sites act independently; that is, there is neither obvious coupling of release sites across an active zone, nor evidence of lateral inhibition^35^. By contrast, at low calcium concentrations, adjacent vesicles tend to fuse together, possibly via a common calcium microdomain.

### Docking is not a stable state

Docked vesicles are often referred to as release-ready vesicles^36^. Indeed, numbers of docked vesicles were profoundly decreased after stimulation (Fig. 2c, ‘all synapses’). However, the degree of docked vesicle depletion is much more severe than expected from the number of pits we observed. At 1.2 mM Ca^2+^, the median number of docked vesicles decreased from 10 to 6 following stimulation for all synapses (p < 0.001; see Supplementary Fig. 3c-d for number of docked vesicles and pits per 10000 nm^2^ of active zone membrane). Therefore, an average of ∼4 pits should be observed in every active zone, but only 35% of synapses contained exocytic pits - consistent with the percentage of active synapses in these cultures^21^. To match the loss of docked vesicles, among these 35% there would need to be an average of ∼10 vesicles fusions per active zone. However, in the active zones that contained pits, the median number of pits was just 1. Likewise, at 2 mM Ca^2+^, docked vesicles decreased from 12 to 8 at 2 mM (p < 0.001), the median number of pits was 2 per pit-containing active zone. At 4 mM Ca^2+^ docked vesicles decreased from 11 to 6 (p < 0.001), the median number of pits was 2 per pit-containing active zone. Increasing the external calcium concentration did not augment the percentage of active zones that respond to an action potential (∼35 %). Thus, we either missed a massive number of fusions (>80% to account for the loss of docked vesicles) or observed activity-dependent undocking of synaptic vesicles^17^.

Interestingly, synapses that did not have pits also exhibited a profound and roughly equal depletion of docked vesicles (Fig. 2c ‘w/o pits’ vs ‘w/pits’). One could imagine that non-responding synapses were just smaller and initially had fewer docked vesicles^37^. However, the active zone size was comparable between those with and without pits (Supplementary Fig. 3e). The absence of pits in these synapses suggests that these synapses are inactive, and the loss of docking at these synapses is not just the result of fusions that we failed to detect. These data imply that docking is not a stable state and that vesicles can stay docked, fuse, or potentially undock upon stimulation.

### Fusing vesicles at 11 ms represent asynchronous release

To determine if vesicles continue to fuse after the 5 ms time point and how these release sites are reoccupied on a short time scale, we performed morphometry on synaptic profiles frozen 5, 8, 11, and 14 ms after an action potential (Fig. 3a-f, Supplementary Fig. 4a-b; 1.2 mM Ca^2+^, 37 °C). Pits peaked at 5 and 8 ms then declined to baseline by 14 ms (Fig. 3g; pits per profile: no stim 0.02, 5 ms 0.21, 8 ms 0.19, 11 ms 0.09, 14 ms 0.03; see Supplementary Fig. 4g for number of pits per 100 nm of active zone). The depth of pits at 5 ms was variable (Fig. 3h; median = 16.2 nm, interquartile range: 13.2 to 22.7 nm), suggesting that some pits have collapsed by this time. Unexpectedly, pits at 11 ms were slightly deeper than those at 5 ms (Fig. 3h; median at 5 ms 16.2 nm; at 11 ms 21.7 nm; p = 0.05). The presence of deep pits suggests that fusion of these vesicles may have initiated later, and may therefore represent asynchronous release^38^.

**Figure 3.**
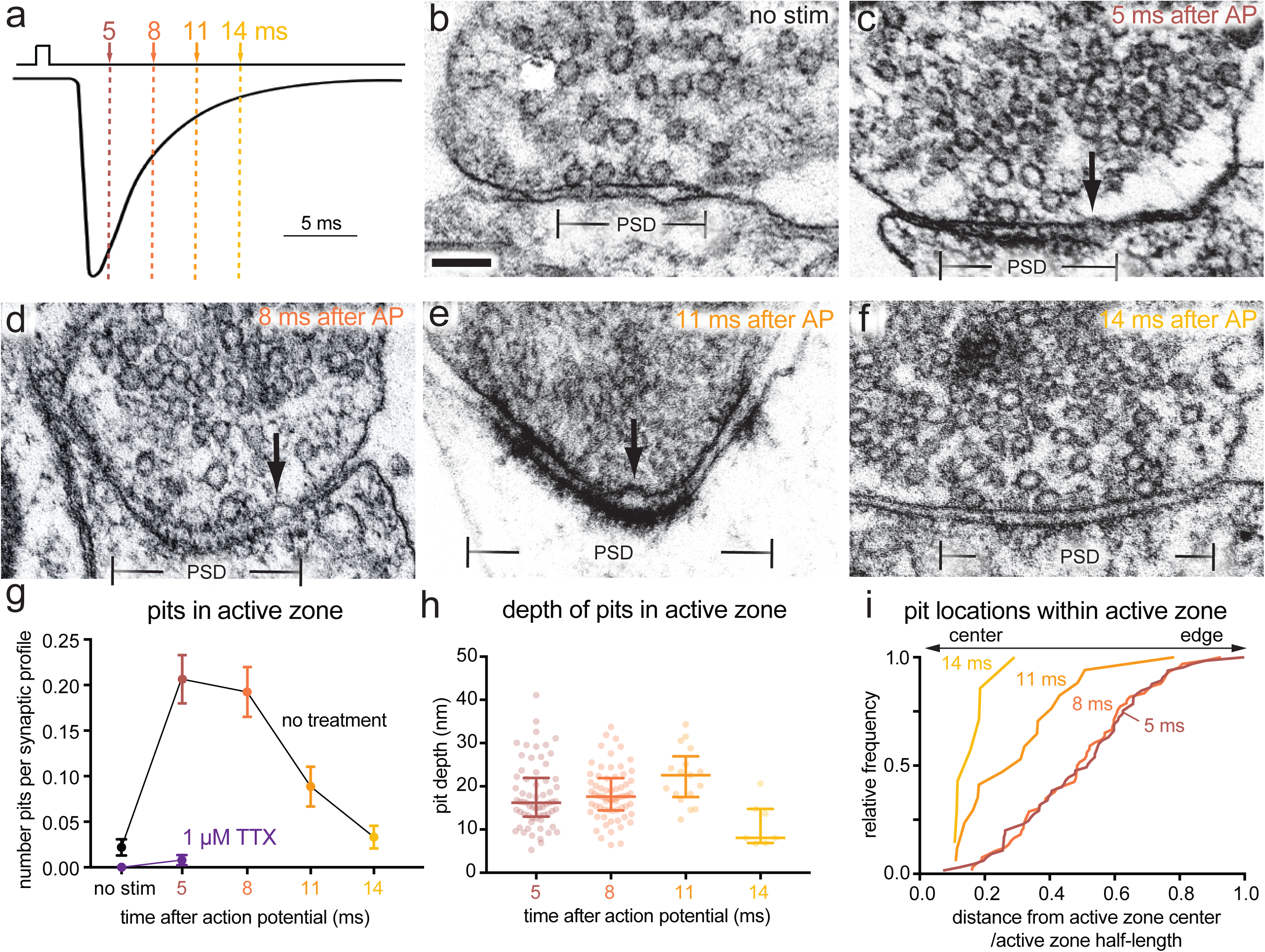
**Vesicle fusion during the first 14 ms after an action potential. a,** Experimental design for stimulation and freezing, showing a diagrammatic excitatory postsynaptic potential for reference (based on ^39^). A 1-ms square pulse is applied to trigger a single action potential, then neurons are frozen at the indicated time points after the beginning of the pulse. **b-f**, Example transmission electron micrographs of synapses from neurons frozen either **b** without stimulation, **c** 5 ms, **d** 8 ms, **e** 11 ms, or **f** 14 ms after stimulation. Arrows indicate pits in the active zone, which are presumed to synaptic vesicles fusing with the plasma membrane. **g**, Number of pits in the active zone per synaptic profile (part of the synapse captured in a 2-D section) in the above conditions, and without stimulation or 5 ms after stimulation in 1 μM tetrodotoxin (TTX, purple); (no stim, n = 274; 5 ms, n = 315; 8 ms, n = 343; 11 ms, n = 192; 14 ms, n = 211; TTX, no stim, n = 121; and TTX, 5 ms, n = 255 synaptic profiles). Numbers of pits with and without stimulation in TTX were not significantly different (p > 0.9). Numbers of pits at 14 ms and without stimulation were not significantly different (p > 0.9). **h**, Depth of pits within the active zone 5 ms (n = 65 pits), 8 ms (n = 66 pits), 11 ms (n = 17 pits), and 14 ms (n = 7 pits) after stimulation. the depth of pits at different time points were all similar (p > 0.05), except for 11 ms and 14 ms, which were significantly different from each other (p = 0.002). **i**, Location within the active zone of the same pits described in **h**. The pits at 11 ms were biased toward the center of the active zone (p = 0.004), while those at 5 ms and 8 ms were not biased toward the center or the edge (p > 0.9 in both cases). Scale bar: 100 nm. PSD: post-synaptic density. AP: action potential. Error bars in **g** indicate standard error of the mean; error bars in **h** and **i** indicate median and interquartile range. All data are from two experiments from separate cultures frozen on different days, except for the data from TTX treatment without stimulation, which are from a single experiment, and data from 5 and 8 ms, which are from three experiments. Numbers of pits in **g**, locations of pits in **h**, and heights of pits in **i** were compared using Kruskal-Wallis tests with full pairwise comparisons by post-hoc Dunn’s multiple comparisons tests. Bias of pit locations toward the center or edge of the active zone was tested by comparing each group to a theoretical median of 0.5 using one-sample two-tailed Wilcoxon signed-rank tests; Bonferroni correction was applied to all p-values from multiple-sample and one-sample tests to account for these extra comparisons. See Supplementary Data Table 1 for full pairwise comparisons and summary statistics. See Supplementary Data Table 2 for summary statistics of pit counts for each experimental replicate.

To specifically test for asynchronous fusion, we assayed exocytosis in the presence of the slow calcium chelator EGTA-AM (25 µM). Intracellular EGTA has a minor effect on synchronous release at most synapses^39^ because the delay between calcium influx and vesicle fusion is less than a millisecond^40^. By contrast, it abolishes slower, asynchronous release^41^. In controls treated with DMSO, pits were apparent in active zone profiles at 5 and 11 ms (Fig. 4a; pits per synaptic profile: at 5 ms 0.16 pits; at 11 ms 0.14 pits; see Supplementary Fig. 5a for more micrographs, 5c for active zone sizes, and 5e for number of pits per 100 nm of active zone). Treatment with 25 μM EGTA-AM had no effect at 5 ms, but eliminated fusion events at 11 ms (Fig. 4b-c; pits per synaptic profile: 5 ms 0.18 pits; 11 ms 0.04, pits, p<0.001; see Supplementary Fig. 5b for more micrographs). Thus, collapse of newly-fused vesicles must be rapid – less than 11 ms; the speed of collapse is thus faster than our previously calculated time constant of 20 ms^7^. These data demonstrate that fusion events observed at 5 and 11 ms represent synchronous and asynchronous release, respectively.

**Figure 4.**
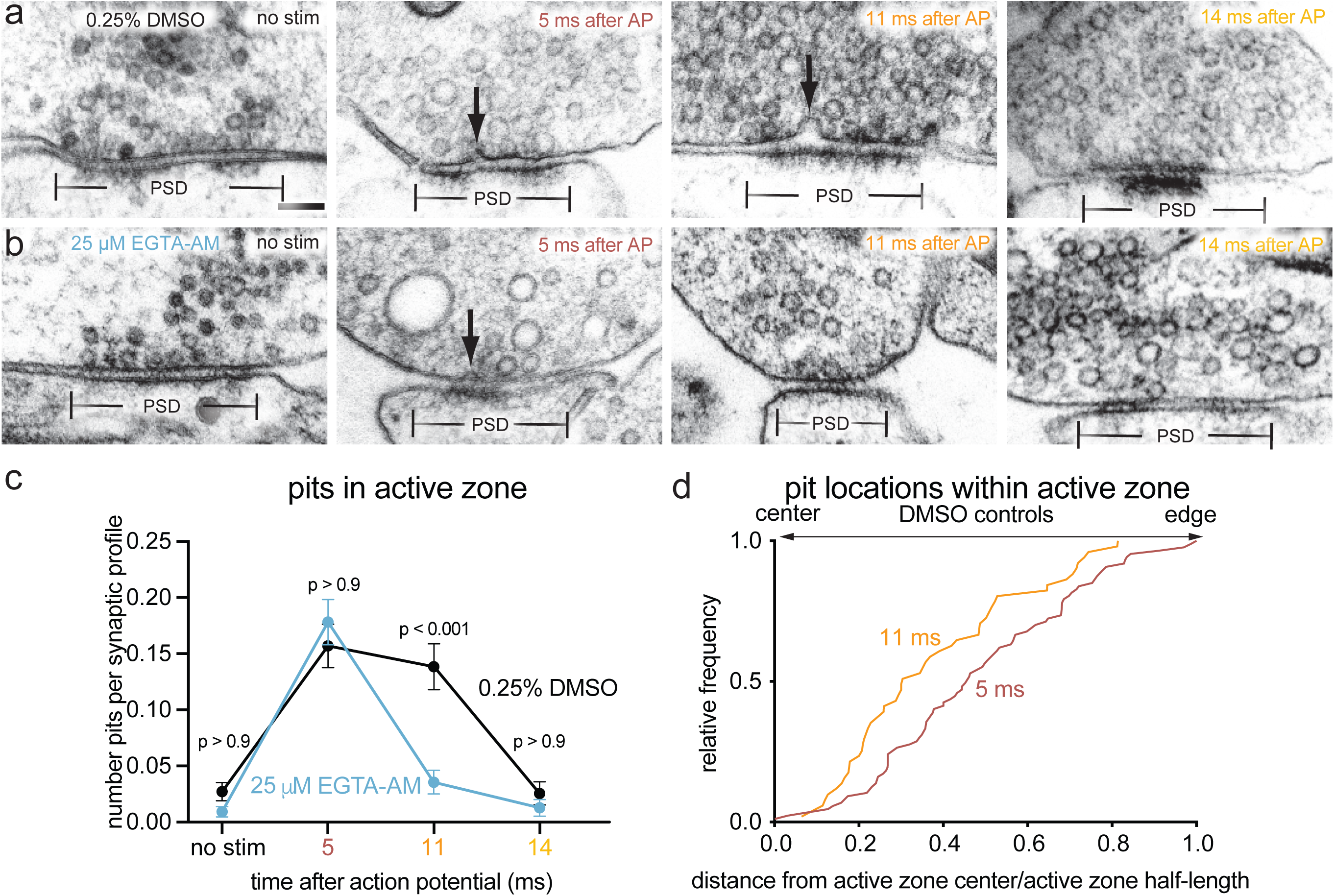
**Fusions captured at 5 and 11 ms after an action potential represent synchronous and asynchronous release. a-b**, Example transmission electron micrographs of synapses from neurons pre-treated with **a** 0.25% DMSO or **b** 25 μM EGTA-AM and frozen either without stimulation, 5 ms after stimulation, 11 ms after stimulation, or 14 ms after stimulation. Arrows indicate pits in the active zone, which are presumed to be synaptic vesicles fusing with the plasma membrane. **c**, Number of pits in the active zone per synaptic profile (part of the synapse captured in a 2-D section) in the above conditions. p-values are from comparisons between EGTA- (no stim, n = 430; 5 ms, n = 421; 11 ms, n = 365; 14 ms, n = 236 synaptic profiles) and DMSO-treated (no stim, n = 405; 5 ms, n = 465; 11 ms, n = 318; 14 ms, n = 235 synaptic profiles) samples frozen at the same time point. Numbers of pits at 11 ms and without stimulation in EGTA-AM-treated samples were not significantly different (p > 0.9). **d**, Locations of pits within the active zone 5 ms (n = 87 pits) and 11 ms (n = 51 pits) after stimulation from neurons pre-treated with 0.25% DMSO. Pits at 11 ms were significantly biased toward the center of the active zone (p < 0.001), while those at 5 ms were not biased toward the center or the edge (p > 0.9). Scale bar: 100 nm. PSD: post-synaptic density. AP: action potential. Error bars in **c** indicate standard error of the mean. All data are from 2-4 experiments from separate cultures frozen on different days. Numbers of pits in **c** were compared using a Kruskal-Wallis test with full pairwise comparisons by post-hoc Dunn’s multiple comparisons test (only comparisons between the same time point with and without EGTA-AM are shown). Locations of pits in **d** were compared using a two-tailed Wilcoxon rank-sum test. Bias of pit locations toward the center or edge of the active zone was tested by comparing each group to a theoretical median of 0.5 using one-sample two-tailed Wilcoxon signed-rank tests; Bonferroni correction was applied to all p-values from two-sample and one-sample tests to account for these extra comparisons. See Supplementary Data Table 1 for full pairwise comparisons summary statistics. See Supplementary Data Table 2 for summary statistics of pit counts for each experimental replicate.

### Asynchronous fusion is concentrated at the center of the active zone

Vesicle fusions occurring during synchronous release were found throughout the active zone, with a slight depletion at the center, in 3D reconstructions of synapses (Fig. 2b). Docked vesicles were generally found throughout the active zone without bias toward the center or edge (Supplementary Table 1 for details; Fig. 2b). Following stimulation, the distribution of docked vesicles within the active zone was unchanged (Fig. 2b; p > 0.1 for each). However, pits were slightly less abundant at the center at all calcium concentrations (Fig. 2b), suggesting that vesicles at the center are initially less fusion-competent.

In single profiles, a lack of bias was also observed during synchronous release; pits and docked vesicles at 5 and 8 ms were not biased toward the center or edge of the active zone (Fig. 3i, 4d, and Supplementary Fig. 4c; 5 ms and 8 ms, p > 0.4 in all cases). By contrast, pits at 11 ms and 14 ms were found near the center of the active zone more frequently, and these distributions were significantly different from those at 5 and 8 ms (Fig. 3i, p = 0.004 and Fig. 4d, p < 0.001). Together, these data argue that vesicles fuse throughout the active zone during synchronous release, whereas asynchronous release is concentrated near the center of the active zone.

### Vesicles transiently dock after synchronous release

As synaptic vesicles are consumed during synchronous and asynchronous release, new vesicles must be recruited to the active zone. During synchronous fusion, docked vesicles across all synaptic profiles were reduced by ∼40% (Fig. 5a-b; docked vesicles per profile: no stim 1.6 vesicles; 5 ms 0.9 vesicles; 8 ms 1.0 vesicles; p < 0.001; see Fig. 2c for 3D analysis at 5 ms; see Supplementary Fig. 4h for number of docked vesicles per 100 nm of active zone). During this time, the number of vesicles close to the membrane but not docked (between 6-10 nm) increased slightly (Fig. 5c, Supplementary Fig. 5d), possibly reflecting vesicles undocked from the active zone (Fig. 2c) or recruited from the cytoplasm. Such vesicles may provide a pool to replace those consumed by fusion. During asynchronous fusion, docked vesicles were not further depleted despite ongoing fusion, implying that synaptic vesicles are recruited during this process (Fig. 5a, 11 ms time point; 1.0 docked vesicles per synaptic profile; p > 0.9 vs 5 ms and 8 ms; p < 0.001 vs no stim). Strikingly, at 14 ms docked vesicles were fully restored to pre-stimulus levels (Fig. 5a; 1.4 docked vesicles per profile, p > 0.9 vs no stim).

**Figure 5.**
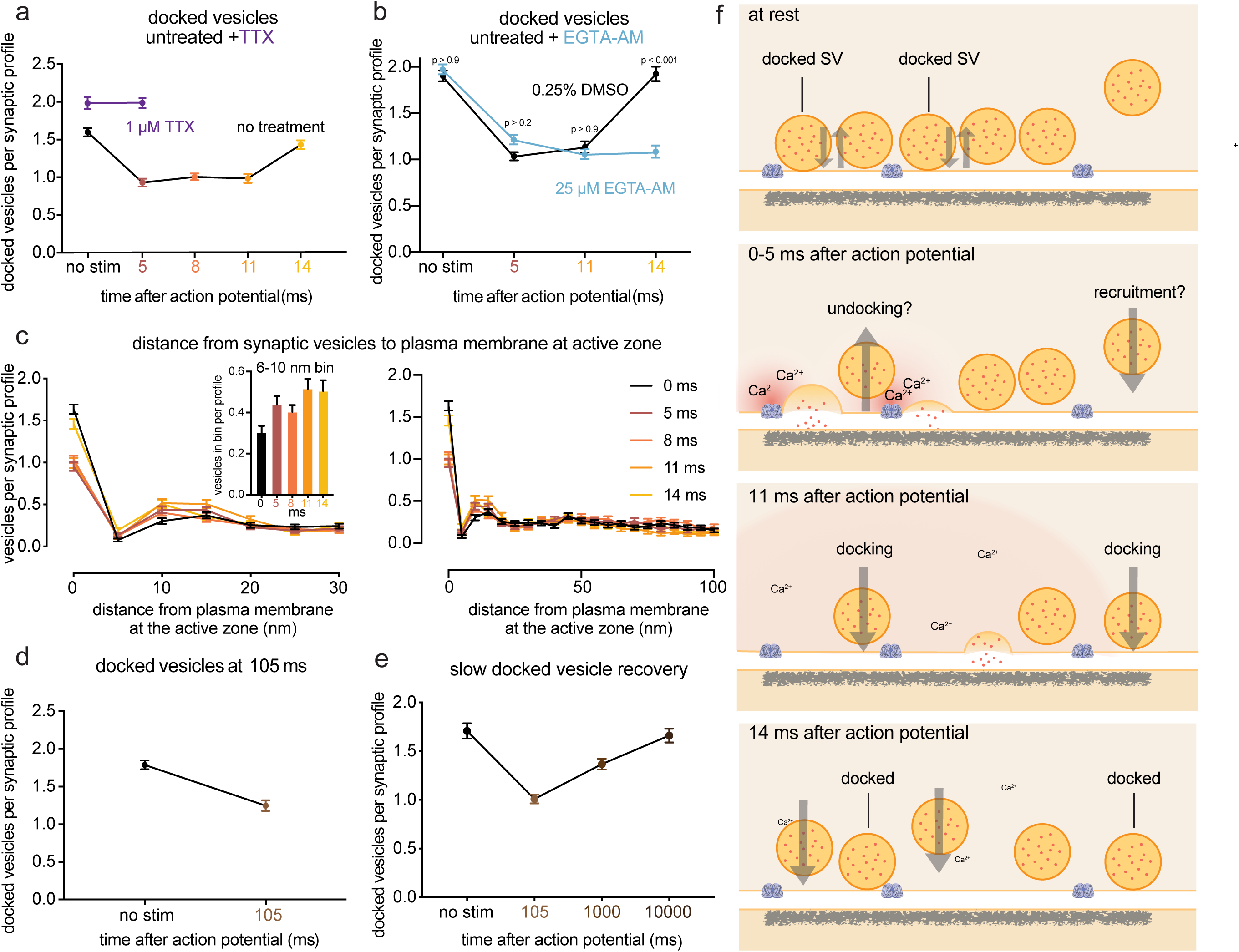
**Transient docking refills the docked vesicle pool within milliseconds. a**, Number of docked vesicles per synaptic profile (part of the synapse captured in a 2D section) from the same experiments and synaptic profiles as in Figure 3. Number of docked vesicles at 14 ms was not significantly different from the no-stimulation control (p > 0.9). **b**, Same as **a**, except from the experiments in Figure 4. Number of docked vesicles at 14 ms in the DMSO control was not significantly different from the no-stimulation control (p > 0.9), but was significantly different with EGTA-AM treatment (p < 0.001). **c**, Distances of synaptic vesicles from the plasma membrane at the active zone, including both vesicles that were annotated as docked and not docked (inset: zoom-in of the 6-10 nm bin). Distances are binned in 5-nm increments, except for “0”, which indicates vesicles ∼0 nm from the active zone membrane (“5” indicates vesicles 0.1-5 nm from the membrane, “10” indicates 6-10 nm, etc.). The number of vesicles at 0 (docked) was significantly greater in the no-stim control than at all other time points but 14 ms. (see p-values listed for **a**). The only other multiplicity-corrected p-values less than 0.05 at any distance were for 11 ms (p = 0.014) and 14 ms (p = 0.048) at 6-10 nm (shown in zoomed inset). **d**, Number of docked vesicles without stimulation or 105 ms after an action potential (no stim, n = 209; 105 ms, n = 218 synaptic profiles). **e**, Number of docked vesicles without stimulation or 105 ms, 1 s, or 10 s after an action potential with 4 mM extracellular [Ca^2+^] (no stim, n = 205; 105 ms, n = 328; 1 s, n = 313; 10 s, n = 212). Vesicles that appeared to be in contact with the plasma membrane were considered docked. Error bars indicate standard error of the mean. Numbers of docked vesicles in **a** and **b** were compared using a Kruskal-Wallis test with full pairwise comparisons by post-hoc Dunn’s multiple comparisons test; Distances of synaptic vesicles from the active zone in the first 5 bins of data shown in **c** (0-25 nm from active zone) were compared to the no-stim control using a one-way ANOVA with post-hoc Games-Howell’s test, with all pairwise comparisons further multiplicity corrected using the method of Bonferroni to account for the 5 ANOVAs. The number of docked vesicles in **d** were compared using a two-tailed Wilcoxon rank-sum test. The number of docked vesicles in **e** were compared using a Kruskal-Wallis test with post-hoc Dunn’s test. Error bars represent standard error of the mean. See Supplementary Data Table 1 for full pairwise comparisons and summary statistics. See Supplementary Data Table 2 for summary statistics for each experimental replicate. **f**, Schematic of events at the active zone of a hippocampal bouton within the first 15 ms after an action potential. Vesicles close to the active zone are proposed to transit between docked and undocked states, with the on- and off-rates resulting in a certain number of vesicles docked and ready to fuse at any given time. Synchronous fusion, often of multiple vesicles, begins throughout the active zone within hundreds of microseconds, and the vesicles finish collapsing into the plasma membrane by 11 ms (note that the high local calcium shown only lasts ∼100 microseconds). Between 5 and 11 ms, residual calcium triggers asynchronous fusion, preferentially toward the center of the active zone. Although shown here as taking place in the same active zone, the degree to which synchronous and asynchronous release may occur at the same active zone after a single action potential is unknown. By 14 ms, the vesicles from the peak of asynchronous fusion, which can continue for tens to hundreds of milliseconds, have fully collapsed into the plasma membrane, and new docked vesicles, which may start to be recruited in less than 10 ms, have fully replaced the vesicles used for fusion. These vesicles then undock or fuse within 100 ms. Whether these new vesicles dock at the same sites vacated by the fused vesicles, and whether newly docked vesicles contributed to synchronous and asynchronous fusion during the first 11 ms, remains to be tested.

Replacement of many forms of release-ready vesicles are known to depend on calcium^15, 42, 43^. We tested whether this ultrafast docking was sensitive to intracellular calcium chelation. Cells were treated with EGTA-AM for 30 min, stimulated and then frozen. EGTA treatment had no effect on the number of docked vesicles in unstimulated samples. Nor did EGTA alter the number of vesicles docked at 5 ms or 11 ms compared to the control (Fig. 5b, no-stim control 2.0; 5 ms time point 1.2; 11 ms time point 1.1 docked vesicles per synaptic profile; p > 0.2 vs DMSO control for each; see Supplementary Fig. 5f for number of docked vesicles per 100 nm of active zone). However, at 14 ms docked vesicles no longer recovered to baseline (Fig. 5b, 14 ms time point 1.1 docked vesicles per synaptic profile; p > 0.9 vs 5 ms and 11 ms; p < 0.001 vs no stim; p < 0.001 vs DMSO 14 ms). These data indicate that the fast recovery of docked vesicles occurring during vesicle fusion is calcium-dependent.

We previously observed that docked vesicle replenishment was slow: docked vesicles were depleted by 50 ms and returned to the baseline by 10 s with a time constant of 3.8 s^7^. Similarly, using zap-and-freeze, docked vesicles were reduced by 30% at ∼100 ms (Fig. 5d; docked vesicles per synaptic profile: no stim 1.8 vesicles; 105 ms 1.2 docked vesicles; p < 0.001). Docked vesicle levels were still 25% lower than unstimulated samples 1 s after stimulation; docking was fully recovered by 10 s (Fig. 5e; docked vesicles per synaptic profile: no stim 1.71; 100 ms 1.0; 1 s 1.37; 10 s 1.66; p < 0.001 between no stim and 105 ms. p = 0.002 between no stim and 1 s, p > 0.9 between no stim and 10 s). Therefore, the fast, calcium-dependent replenishment of docked vesicles observed at 14 ms is temporary and appears to be lost within 50 ms; transient docking is followed by slower docking process that requires 3-4 sec. Transient docking^9^ could provide fusion-competent vesicles for asynchronous release^16^ and counteract synaptic depression during trains of stimuli^9^.

## Discussion

We characterized docking and exocytosis of synaptic vesicles at hippocampal synapses in ultrastructural detail (Fig. 5f). Cultured neurons were subjected to a single stimulus, frozen at defined time points, and processed for electron microscopy. We performed morphometry on 6071 single synaptic profiles and 389 fully reconstructed active zones. Although ∼65% of synapses do not respond to a single action potential, multivesicular release is common at the 35% of synapses that respond to an action potential. Fusion sites are dispersed throughout the active zone, and vesicles fully collapse into the plasma membrane. Asynchronous fusion, defined as delayed events that are sensitive to the slow calcium buffer EGTA, occurs preferentially near the center of the active zone. By 14 ms, fused vesicles are fully replaced by new transiently docked vesicles, which return to the undocked state in less than 100 ms. These findings have implications for synaptic failure, multivesicular release, the spatial organization of release sites, and their refilling during short-term plasticity.

Roughly 65% of synapses in cultured hippocampal neurons do not respond to a single action potential as determined by the presence of fusion pits after stimulation. This failure rate is consistent with previous experiments in which we stimulated using channelrhodopsin rather than using electrical stimulation and assayed activity by endocytosis rather than by vesicle fusion^7, 21^. This failure rate in itself is not unusual; the fraction of synapses that respond to a single action potential *in vivo* is low for many neurons of the central nervous system^44^. A 65% failure for individual synapses is consistent with the mean failure rate determined using a minimal stimulation protocol^27^ or by use-dependent block of NMDA glutamate receptors^28, 29^. For example, in autaptic cultures 65% of synapses were found to have a low probability of release and 35% a high probability of release (C. Rosenmund, personal communication). Given that we stimulated only once, it is possible that all synapses have a uniform release probability of 0.35 and release is stochastic; that is, different synapses would be recruited with every stimulation. However, a large fraction of synapses in our cultures appear to be completely silent: First, increased calcium does not recruit more synapses: only ∼35% of synapses exhibited vesicle fusions at 1.2 mM, 2 mM or 4 mM calcium. Second, in previous experiments, we used dynasore to block dynamin during multiple stimulations^39^. Only slightly more than 30% of the synapses exhibited trapped endocytic structures when stimulated 100 times. We conclude that most synapses in our cultures are not active even during high-frequency stimulation.

How many release sites are there within an active zone as determined by docking sites? The median number of docked vesicles in our experiments was ∼10 per synapse. Previous studies have observed means of 5-10 docked vesicles at hippocampal synapses^3, 4, 37, 45, 46^. The number of release sites at cultured hippocampal synapses determined using various fluorescence techniques was also 5-10^32, 47–49^. Similarly, the docking and priming protein Munc13 was typically found in 6 clusters per synapse^32^ (range 0-17). Thus, the number of docked vesicles is similar to the number of release sites.

The presence of multiple vesicles docked at a synapse alone does not imply that multiple vesicles can fuse at an active zone. In fact, it was long thought that only one vesicle could fuse in response to an action potential^24, 50^. These studies argued that responses at synapses are mostly, or even exclusively, uniquantal, despite the presence of multiple docked vesicles. One mechanism put forward for univesicular release is that fusion of a vesicle can alter the active zone by “lateral inhibition” and thereby block further fusions^35, 51^. For proponents of univesicular release, examples of recordings of multivesicular events were dismissed as being caused by multiple active zones impinging on the cell. Proponents of multiquantal release at single active zones argued that observations of uniquantal events were due to saturation of the postsynaptic receptor field, and multiquantal release could be observed under circumstances in which saturation could be avoided ^52, 14, 15, 16, 20^. By reconstructing synapses from serial sections immediately after a single action potential, we were able to capture multiple vesicles fusing in a single active zone. At 4 mM calcium, we observed up to 11 vesicles fusing in a single active zone. The probability of fusion at a release site appears to be low even in elevated calcium, but because active zones have ∼10 docking sites, multiple vesicles can be consumed by a single action potential.

Importantly, fusing vesicles tended to be close (<100 nm) to one another at low calcium concentrations, and were in fact often adjacent. Adjacent fusions can also observed during spontaneous activity: in a previous study, 20% of synaptic profiles exhibiting spontaneous fusions comprised adjacent fusions, suggesting that fusing vesicles are coupled even in the absence of stimulation^46^. It is likely that coupled fusion is being driven by an active calcium channel or calcium microdomain that acts on locally docked vesicles^40, 56, 57^.

In contrast to the microdomains that drive synchronous release, the residual calcium that triggers asynchronous release is more broadly distributed and longer-lasting^38, 58^. This implies that there would be no spatial specificity for asynchronous fusion. However, we found that asynchronous release occurs preferentially near the center of the active zone, whereas synchronous fusion is evenly distributed across the active zone. Several molecules, including VAMP4^59^, Synaptotagmin-7^60^, SNAP23^61^, and Doc2^62^, have been implicated in asynchronous release, and these molecules could target vesicles to release sites near the center of an active zone. Alternatively, the locations of voltage-gated calcium channel clusters within an active zone may account for this spatial arrangement. In both *Caenorhabditis elegans*^63^ and *Drosophila melanogaster*^64^ neuromuscular junctions, different isoforms of Unc13 position vesicles at different distances from the dense projection, where calcium channels reside^65^. Based on phenotypes from isoform-specific knockouts, these clusters were proposed to form independent release sites for fast and slow phases of neurotransmission. In conclusion, while the molecular mechanism remains uncertain, synchronous and asynchronous release are concentrated in different regions of the active zone.

A profound decrease in docking was observed after stimulation. The fusions we observed can only account for ∼30% of the decrease in docking. Moreover, docking is also reduced in synapses with no visible fusions. The loss of docked vesicles is accompanied by a slight increase in vesicles 6-10 nm from plasma membrane, suggesting that these vesicles may still be tethered to the membrane by a loosely assembled SNARE complex, synaptotagmin, or Munc13^4^. However, it is equally possible that more vesicles are recruited to this region from the cytoplasm. Furthermore, the increase in vesicles 6-10 nm from the plasma membrane also cannot fully account for the massive loss of docked vesicles, leaving their fate uncertain. Therefore, we conclude that either there is far more exocytosis than anticipated (∼40% of docked vesicles across all synapses fusing after a single action potential), vesicles become undocked after a single action potential, or some combination of the two.

At 14 ms after the stimulation, docking levels are fully restored to pre-stimulus levels. But then by 100 ms after stimulation, docking is again reduced to the levels observed immediately after stimulation– thus the docking that occurs 10-14 ms after the action potential is transient. We did not observe transient docking in our previous ‘flash-and-freeze’ experiments^7^, likely because the generation and timing of action potentials using channelrhodopsin is unreliable. However, the more prolonged reduction in docked vesicles observed here at 100 ms and 1 s is consistent with our previous results^7^. Full and stable restoration of docking was not restored until 3-10 s^7^ after stimulation, consistent with the recovery of the physiological readily-releasable pool^13^. Thus, there is a rapid docking of vesicles after stimulation, but this docking is only transient.

What purpose could fast, but ephemeral, vesicle recruitment serve? Quite likely it is to maintain robust synaptic transmission during trains of stimuli. Recent electrophysiological studies of a cerebellar ‘simple synapse’ comprised of a single active zone indicate that an undocked population of vesicles may occupy a ‘replacement site’^15, 16^, possibly corresponding to our 10 nm pool. Based on modeling, vesicles in this pool are rapidly mobilized to dock at a release site, peaking ∼10 ms after a stimulus. However, these docked vesicles become undocked and return to the replacement site in the 100 ms following the action potential. Transient docking is likely mediated, at least in part, by the calcium sensor Synaptotagmin-1^9^. When Synaptotagmin-1’s membrane-binding residues were mutated, vesicle docking was reduced by 30-50%. Docking was restored by an action potential, but had declined after 100 ms, consistent with the time course of docking that we observed. Our data demonstrate that transient docking is not just a quirk of Synaptotagmin-1 mutants. Moreover, vesicles may undock before transiently redocking.

In summary, we have characterized the ultrastructure of a synapse during the first 14 ms after an action potential using zap-and-freeze electron microscopy. At physiological calcium concentrations, an action potential drives fusion of one or more vesicles, likely via a shared calcium microdomain. It is presumed that such vesicles are docked to the membrane in a “tight-state” as recently proposed^17^. Stimulation is accompanied by a massive reduction of the docked pool, perhaps even in synapses that do not exhibit fusion. One possibility is that calcium drives docked vesicles into an undocked state, most likely by binding to a protein such as Munc13 or Synaptotagmin-1, or possibly to a lipid such as PIP2. Alternatively, vesicles could be in a dynamic equilibrium between fusion-competent (docked) and -incompetent states (undocked) at steady state near the active zone membrane, and only those that are tightly docked coincident with calcium influx would fuse. These undocked vesicles would still be associated with release sites but are tethered ∼10 nm from the membrane. Such vesicles are proposed to exist in a “loose-state” with SNAREs, Synaptotagmin-1, and Munc13 still engaged^66^. Between 8 and 14 ms, vesicles dock to the membrane in a calcium-dependent manner, perhaps driven by Synaptotagmin-1 or the calcium sensor for facilitation, Synaptotagmin-7^67^. Docking is occurring at the same time as vesicles are undergoing asynchronous fusion and may represent vesicles undergoing ‘2-step’ release^16^. Docking levels are fully restored 14 ms after stimulation; however, this docking is not stable, and declines along with falling calcium levels. This time course is similar to that of paired-pulse facilitation of synaptic transmission, which only lasts for milliseconds^68^. Thus, synaptic vesicles at the active zone exhibit surprisingly lively dynamics between docked and undocked states within milliseconds after an action potential.

## Methods

All animal care was performed according to the National Institutes of Health guidelines for animal research with approval from the Animal Care and Use Committee at the Johns Hopkins University School of Medicine.

### Neuronal cell culture

Cell cultures were prepared on 6-mm sapphire disks (Technotrade), mostly as previously described^7, 21^. Newborn or embryonic day 18 C57/BL6J mice of both sexes were decapitated, followed by dissection of and transfer of brains to ice-cold HBSS. In the case of embryonic mice, heads were stored in HBSS on ice prior to dissection. For high-pressure freezing, neurons were cultured on a feeder layer of astrocytes. For FM dye experiments, astrocytes were grown on 22- mm coverslips for 1 week and placed on top of neurons cultured on sapphire disks with astrocytes facing neurons^69^, with Paraffin dots used as spacers. Astrocyte cultures were established from cortices trypsinized for 20 min at 37 °C with shaking, followed by trituration and seeding on T-75 flasks. Astrocytes were grown in DMEM supplemented with 10% FBS and 0.1% penicillin-streptomycin for 2 weeks, then plated on PDL-coated 6mm sapphire disks atop glass coverslips in 12-well plates at a density of 50,000/well to create a feeder layer. After six days, FUDR was added to stop cell division. The following morning, culture medium was replaced with Neurobasal-A supplemented with 2% B27 and 0.1% penicillin-streptomycin (NB-A full medium, Invitrogen) prior to plating hippocampal neurons. Hippocampi were dissected and incubated in papain with shaking at 37 °C for 30-60 min, then triturated and plated on astrocytes at 50,000 or 75,000 cells/well. Before use, sapphire disks were carbon-coated with a “4” to indicate the side that cells are cultured on. Health of the cells, as indicated by de-adhered processes, floating dead cells, and excessive clumping of cell bodies, was assessed regularly, as well as immediately before experiments. All experiments were performed between 13 and 17 days *in vitro*.

### Electrical field stimulation

The electrical stimulator is manufactured by Leica to be compatible with the Leica ICE high pressure freezer. The middle plate was designed as a circuit board trimmed to the dimensions of a standard Leica ICE high-pressure freezer middle plate. In the middle plate, there is a central 6 mm hole holding the sample sandwiched between two sapphire disks. This central hole was plated with two gold contact surfaces that are used to apply field stimulation to the sample. The standard spacer ring between the sapphire disks are conductive, and is replaced with nonconductive mylar rings of the same dimensions. The voltage to be applied to the sample is provided by a capacitor bank attached to the middle plate. The capacitors are charged just before the sample is loaded into the chamber. The current from the capacitors to the sample is controlled by a phototransistor. In the absence of light, there is no current passed from the capacitors to the sample contacts. In this way, the field stimulation can be activated within the chamber using the standard light stimulation function of the EM ICE.

### FM dye uptake imaging and quantification

For the FM 1-43FX (Invitrogen) uptake assay, we used a modified version of a previously published protocol^70^. Neurons on sapphire disks were first incubated with 30 µM Pitstop 2 (Sigma) in physiological saline (1 mM Ca^2+^) for 2 min. This treatment blocks regeneration of synaptic vesicles from synaptic endosomes^21^, so as to prevent FM dyes from being released during the washing procedure. Following addition of FM dye (5 µg/ml), a sapphire disk was mounted on a middle plate, while another sapphire disk in the same well was left in the solution to ensure that both sapphire disks were incubated in FM dye for the same period of time. After charging the middle plate, 10 pulses of light (1 ms each) were applied at 20 Hz to discharge the capacitor and induce 10 action potentials. Immediately after stimulation, both stimulated and unstimulated specimens were transferred to an 18-mm petri dish containing physiological saline solution (1 mM Ca^2+^). FM dyes bound to the plasma membrane were washed off by passing current across the specimen using a transfer pipet for 1 min. Both samples were then transferred into warm (37 °C) PBS containing 4% paraformaldehyde and fixed for 30 min. After fixation, samples were washed 3x with PBS and immediately imaged on an Olympus IX81 epifluorescence microscope equipped with a Hamamatsu C9100-02 EMCCD camera with mercury lamp illumination through a CFP/YFP filter set (Semrock) and a 60x, NA 1.4 Olympus UIS2 oil-immersion objective. For each condition, seven images were acquired and 20 putative presynaptic terminals quantified, identified by their increased FM labeling relative to the rest of the axon, shape, and size, by manual segmentation in ImageJ, and their total fluorescence intensity measured. Intensity values were then background corrected. All micrographs shown were acquired with the same settings on the microscope and later adjusted in brightness and contrast to the same degree in ImageJ, then rotated and cropped in Adobe Photoshop.

### High-pressure freezing

Cells cultured on sapphire disks were frozen using an EM ICE high-pressure freezer (Leica Microsystems). The freezing apparatus was assembled on a table heated to 37 °C in a climate control box, with all solutions pre-warmed (37 °C). Sapphire disks with neurons were carefully transferred from culture medium to a small culture dish containing physiological saline solution (140 mM NaCl, 2.4 mM KCl, 10 mM HEPES, 10 mM glucose; pH adjusted to 7.3 with NaOH, 300 mOsm). NBQX (3 μM; Tocris) and bicuculline (30 μM; Tocris) were added to the physiological saline solution to block recurrent synaptic activity. CaCl_2_ and MgCl_2_ concentrations were 1.2 mM and 3.8 mM, respectively, except where indicated, in which case the MgCl_2_ concentration was adjusted accordingly (3 mM MgCl_2_ with 2 mM CaCl_2_, 1 mM MgCl_2_ with 4 mM CaCl_2_). Cells were then fitted into the photoelectric middle plate. Filter paper was placed underneath the middle plate to remove all excess liquid. A mylar spacer ring was then placed atop the sapphire disk. To create a “sandwich” of the solution described above, the underside of another sapphire disk was dipped in the solution so that some was held on by surface tension, then placed atop the spacer ring so that excess liquid again dispersed onto the filter paper. For voltage to be applied across the sample, it is essential for all components outside of the sandwich to be dry, so the top of the sapphire and all other components of the setup were gently dried with another piece of filter paper. Finally, a rubber ring was added to hold everything in place. This entire assembly process takes 3-5 min. The assembled middle plate was enclosed in two half cylinders then loaded into the freezing chamber, where the cells were stimulated for 1 ms before freezing at the desired time point, ranging from 5 ms to 105 ms. With this protocol, 10 V/cm is applied for 1 ms across a 6-mm space between the electrodes into which the sapphire disk fits, as confirmed by measurements from Leica. This field stimulation regimen in hippocampal cultures induces a single action potential and only negligibly depolarizes boutons directly^23^.

For EGTA experiments, first half of the media in which cells were grown was removed and set aside. EGTA-AM (Fisher) or DMSO was then added to media to a final concentration of 25 μM and 0.25% DMSO for 15 min to load the cells with EGTA. Cells were washed three times and left in the media that had been set aside for 15 min before freezing in the physiological saline solution described above (treatment protocol adapted from ^39^). For TTX experiments, TTX was added to the freezing solution to a final concentration of 1 μM, such that cells were in TTX for 3-5 min before freezing.

Cooling rates during freezing were between 16,000-18,000 K/sec. Membrane traffic stops at 0 °C, so the point at which the sample reaches this temperature can be considered the true time of freezing. On the EM ICE, we set the stimulation program to produce a 1-ms pulse, followed by a resting period (ranging from 0 ms to 10 s). By default, during the freeze process, the temperature sensor placed just outside the specimen chamber reaches 0 °C, precisely when the resting period of the stimulation program is complete. This causes an extra 5-ms delay in samples reaching 0 °C. Specifically, an additional ∼4 ms is needed for the chamber to freeze (∼3 ms faster than the HPM100) and another 1 ms for neurons to freeze^7^. Thus, a total of 5 ms delay is expected. This 5-ms delay was confirmed by direct measurements made by Leica Microsystems. Previous experiments indicated that this number may be off by ± 1 ms due to the mechanics of the EM ICE^71^. Therefore, specimens were frozen, on average, 5 ms later than the time point programmed into the EM ICE, with relatively little variability. For example, to freeze at 5 or 8 ms, the delay period on the machine was set to “0 ms” or “3 ms”. Thus, the time points indicated (5, 8, 11, 14, and 105 ms) are calculated based on this estimated 5 ms delay from the onset of stimulation.

### Freeze-substitution

After freezing, samples were transferred under liquid nitrogen to an EM AFS2 freeze substitution system at -90 °C (Leica Microsystems). Using pre-cooled tweezers, samples were quickly transferred to anhydrous acetone at -90 °C. After disassembling the freezing apparatus, sapphire disks with cells were quickly moved to cryovials containing 1% glutaraldehyde, 1% osmium tetroxide, and 1% water in anhydrous acetone, which had been stored under liquid nitrogen then moved to the AFS2 immediately before use. The freeze substitution program was as follows: −90 °C for 6-10 hr (adjusted so substitution would finish in the morning), 5 °C h^−1^ to −20 °C, 12 h at -20 °C, and 10 °C h^−1^ to 20 °C.

### Embedding, sectioning, and transmission electron microscopy

Samples in fixatives were washed three times, 10 min each, with anhydrous acetone, then stained *en bloc* with 1% uranyl acetate for 1 hr with shaking. After three washes, samples were left in 30% epon araldite in anhydrous acetone for 3 hr, then 70% epon araldite for 2 hr, both with shaking. Samples were then transferred to caps of polyethylene BEEM capsules (EMS) and left in 90% epon araldite overnight at 4 °C. The next morning, samples were transferred to 100% epon araldite (epon, 6.2 g; araldite, 4.4 g; DDSA, 12.2 g; and BDMA, 0.8 ml) for 1 hr, then again to 100% for 1 hr, and finally transferred to 100% epon araldite and baked at 60 °C for 48 hr.

For single-section imaging, 40-nm sections were cut, while 10-15 serial sections were cut for serial-section 3D reconstructions, 50 nm thick in the first replicate and 40 nm in the second. Sections on single-slot grids coated with 0.7% pioloform were stained with 2.5% uranyl acetate then imaged at 80 kV on the 93,000x setting on a Phillips CM 120 transmission electron microscope equipped with an AMT XR80 camera. In some cases, including all serial-section imaging, the microscopist was blind to the different conditions, while in other cases they were not. To limit bias, synapses were found by bidirectional raster scanning along the section at 93,000x, which makes it difficult to “pick” certain synapses, as a synapse usually takes up most of this field of view. Synapses were identified by a vesicle-filled presynaptic bouton and a postsynaptic density. Postsynaptic densities are often subtle in our samples, but synaptic clefts were also identifiable by 1) their characteristic width, 2) the apposed membranes following each other closely, and 3) vesicles near the presynaptic active zone. Only synapses with prominent post-synaptic densities were imaged for serial-sectioning reconstructions. 125-150 micrographs per sample of anything that appeared to be a synapse were taken without close examination. For serial sectioning, at least 30 synapses per sample were imaged.

### Electron microscopy image analysis

Images were annotated blind but not randomized in the initial time course experiments (first replicate of data shown in Figure 3) and the first replicate of the serial-sectioning data in Figure 2. For all other data, all the images from a single experiment were randomized for analysis as a single pool using a custom R (R Development Team) script. Only after this randomization were images excluded from analysis, either because they appeared to not contain a *bona fide* synapse or the morphology was too poor for reliable annotation. This usually meant ∼100 synapses per sample were analyzed for single sections. In some cases, membranes had low contrast against the cytoplasm, due mostly to good preservation of proteins in these tissues. These images are annotated after adjusting the contrast in ImageJ. The plasma membrane, active zone, docked synaptic vesicles, synaptic vesicles close to the active zone, and pits (putative fusion events) were annotated in ImageJ using a custom plugin. The active zone was identified as the region of the presynaptic plasma membrane with the features described above for identifying a synapse. Docked vesicles were identified by their membrane appearing to be in contact with the plasma membrane at the active zone (0 nm from the plasma membrane), that is, there are no lighter pixels between the membranes. When comparing data, note that ‘docking’ is more narrowly defined in these data than in Imig et al.^36^ (0-2 nm) and Chang et al.^9^ (<5nm), and is the definition of docking that we have used in previous publications^5, 7, 8^. Vesicles that were not manually annotated as docked, but were 0 nm away from the active zone plasma membrane, were automatically counted as docked when segmentation was quantitated (see below) for data sets counting the number of docked vesicles. Vesicles annotated as docked were automatically placed in the 0 nm bin of vesicle distances from the plasma membrane. Pits were identified as smooth curvature (not mirrored by the postsynaptic membrane) in an otherwise straight membrane. These pits are considered exocytic, as endocytic pits do not normally appear until 50 ms after an action potential^7^, fluid phase markers are not internalized until ∼100 ms^7^, and ferritin-positive vesicles are not found near the active zone membrane until ∼10 s after stimulation^21^. Pits outside the active zone are considered endocytic or membrane ruffles, as this is the primary site for ultrafast endocytosis^7^.Under these criteria, we could miss or over-annotate vesicles and pits. To minimize the bias and maintain consistency, all image segmentation, still in the form of randomized files, was thoroughly checked by a second member of the lab. For serial-section data, active zones with multiple pits were re-evaluated *post hoc* after unblinding to make sure they are not halves of the same structure. However, no corrections were made for synaptic vesicles since vesicles are much more abundant and the same criteria were used to annotate them in all conditions. A similar amount of overestimate is expected in this case. Features were then quantitated using custom MATLAB (MathWorks) scripts.

Location of pits and docked vesicles within the active zone from single sections was calculated from the distance from the center of the pit to the center and the edge of the active zone in 2D. Distance from the center was normalized by dividing the distance to the edge by the half-width of the active zone. For 3D data, the distance to the center of the active zone was calculated from serial sections. First, the location in 2D was calculated as above. Then, the 3D distance was calculated to the center of the active zone in the middle section of the series using the Pythagorean theorem with assumption that each section is the same thickness and the center of the active zone aligns in each image. Locations in 3D data were further corrected to be the density of vesicles/pits at each distance from the center of the active zone. This is because the total area for objects to be located in increases with increasing distance from the center of a roughly circular object (for example, randomly distributed objects within a circular active zone would have a median distance from the center of 0.66, giving the impression that they are biased toward the edge: after calculating the density, this value would be 0.5). To calculate density of vesicles/pits from the center to the edge in 3D reconstructions, the radial position of each vesicle/pit was converted to the fractional area of a circle bounded by that radius. In the case of a unit circle (distance from center to edge is by definition 1 data normalized to the size of the active zone), this is simply the square of the original normalized distance to the center. Distance between pits and docked vesicles in different sections was approximated in a similar manner, where the edges of the hypothetical triangle are 1) the difference of the distances between each pit to the center of the active zone in each section and 2) the distance between the sections, again assuming a thickness of 50 nm.

Example micrographs shown were adjusted in brightness and contrast to different degrees (depending on the varying brightness and contrast of the raw images), rotated, and cropped in Adobe Photoshop.

### Statistical analysis

All data shown are pooled from multiple experiments; see Supplemental Table 2 for summary data for each replicate. All data were initially examined on a per-experiment basis (with all freezing done on the same day and all segmentation done in a single randomized batch); none of the pooled data show any result that was not found in each replicate individually. We did not predetermine sample sizes using power analysis, but based them (N = 2-3, n > 200) on our prior experience with flash-and-freeze data^7, 8, 21^. An alpha of 0.05 was used for statistical hypothesis testing. All data were tested for normality by D’Agostino-Pearson omnibus test to determine whether parametric or nonparametric methods should be used. Comparisons between two groups were performed using a two-tailed Welch two-sample t-test or Wilcoxon rank-sum test. Comparisons between multiple groups followed by full pairwise comparisons were performed using one-way analysis of variance (ANOVA) followed by Tukey’s HSD test or Kruskal-Wallis test followed by Dunn’s multiple comparisons test. Differences in the number of active zones containing at least one pit from active zone reconstructions in Figure 2a were assessed using a chi-square test. For testing whether locations of pits were biased toward the center or edge of the active zone, a two-tailed one-sample t-test or Wilcoxon rank-sum test with a theoretical median of 0.5 was used (each of these p-values, as well as that of the comparisons between pit locations in different samples, were accordingly corrected for multiplicity using Bonferroni’s method). All statistical analyses were performed and all graphs created in Graphpad Prism 7.

### Life Sciences Reporting Summary

More details on experimental procedures, materials, and statistics are available in the Life Sciences Reporting Summary.

## Data and code availability

The data underlying this work, as well as custom R and MATLAB scripts, are available upon request.

## Acknowledgements

We are indebted to Shuo Li, Quan Gan, Kie Imoto, Delgermaa Lubsanjav, Chengxiu Zhang, and Sebastian Markert, for cell culture, help with freezing, and stimulating discussions. We also thank Mike Delanoy and Barbara Smith for technical assistance with electron microscopy and Kathleen T DiNapoli for developing R code to randomize images. We thank Paul Wurzinger and Cveta Tomova at Leica for design and manufacture of middle plate, Melissa A Herman for initial tests of using a capacitor for field stimulation, Hana Goldschmidt for help validating the stimulation device using pHluorin imaging, and Nathan Livingston for help with voltage imaging. We also thank the Marine Biological laboratory and their Neurobiology course for supporting the initial set of experiments. S.W. and this work were supported by start-up funds from the Johns Hopkins University School of Medicine, Johns Hopkins Discovery funds, and the National Science Foundation (1727260), the National Institutes of Health (1DP2 NS111133-01 and 1R01 NS105810-01A1) awarded to S.W. S.W. is an Alfred P. Sloan fellow, McKnight Foundation Scholar, and Klingenstein and Simons Foundation scholar. E.M.J. is an Investigator of the Howard Hughes Medical Institute. G.F.K. was supported by a grant from the National Institutes of Health to the Biochemistry, Cellular and Molecular Biology program of the Johns Hopkins University School of Medicine (T32 GM007445) and is a National Science Foundation Graduate Research Fellow (2016217537). The EM ICE high-pressure freezer was purchased partly with funds from an equipment grant from the National Institutes of Health (S10RR026445) awarded to Scot C Kuo.

## Author Contributions

M.W.D., S.W., and E.M.J. conceived the zap-and-freeze technique. G.F.K. and S.W. designed the experiments and analyzed the data. S.W., G.F.K., and E.M.J wrote the manuscript. G.F.K. performed all freezing experiments and single-section electron microscopy sample preparation, imaging, and analysis, and FM dye uptake experiments, with technical assistance from S.W, with the exception of the first replicate of the 105 ms time point experiment, which was performed by S.W, and the 1 and 10 s time point experiments, which were performed by S.R. M.C. and G.F.K. performed the serial sectioning 3D reconstruction electron microscopy imaging and analysis. S.W. and M.C. developed MATLAB code for image analysis. K.P.A., E.H., K.L., S.R., and T.V. performed pilot zap-and-freeze experiments and electron microscopy sample preparation, imaging, and analysis. M.W.D. designed the prototype zap-and-freeze stimulation device. S.W. funded and oversaw the research.

---

**Supplementary Figure 1.**
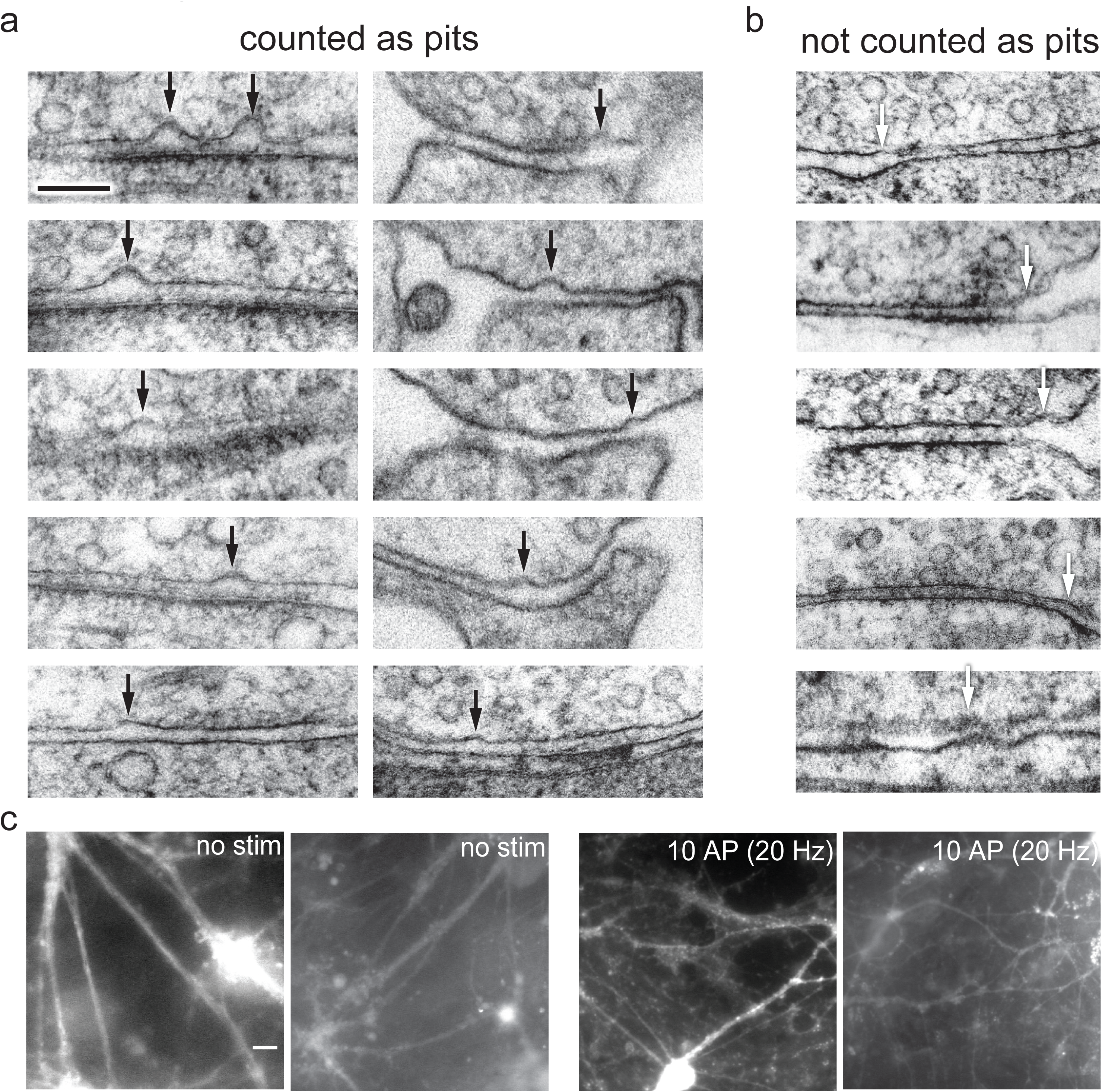
**Examples of pits in the active zone compared to pits outside the active zone or features not quantified as pits. a**, Examples of pits in the active zone 5 ms after stimulation, indicated by black arrows. **b**, Examples of features not counted as pits, for instance because the curvature in the presynaptic membrane is mirrored by the postsynaptic membrane, indicated by white arrows. Scale bar: 100 nm **c**, Full fields-of-view of micrographs from the FM dye experiment described in Figure 4b. Scale bar: 20 microns.

**Supplementary Figure 2.**
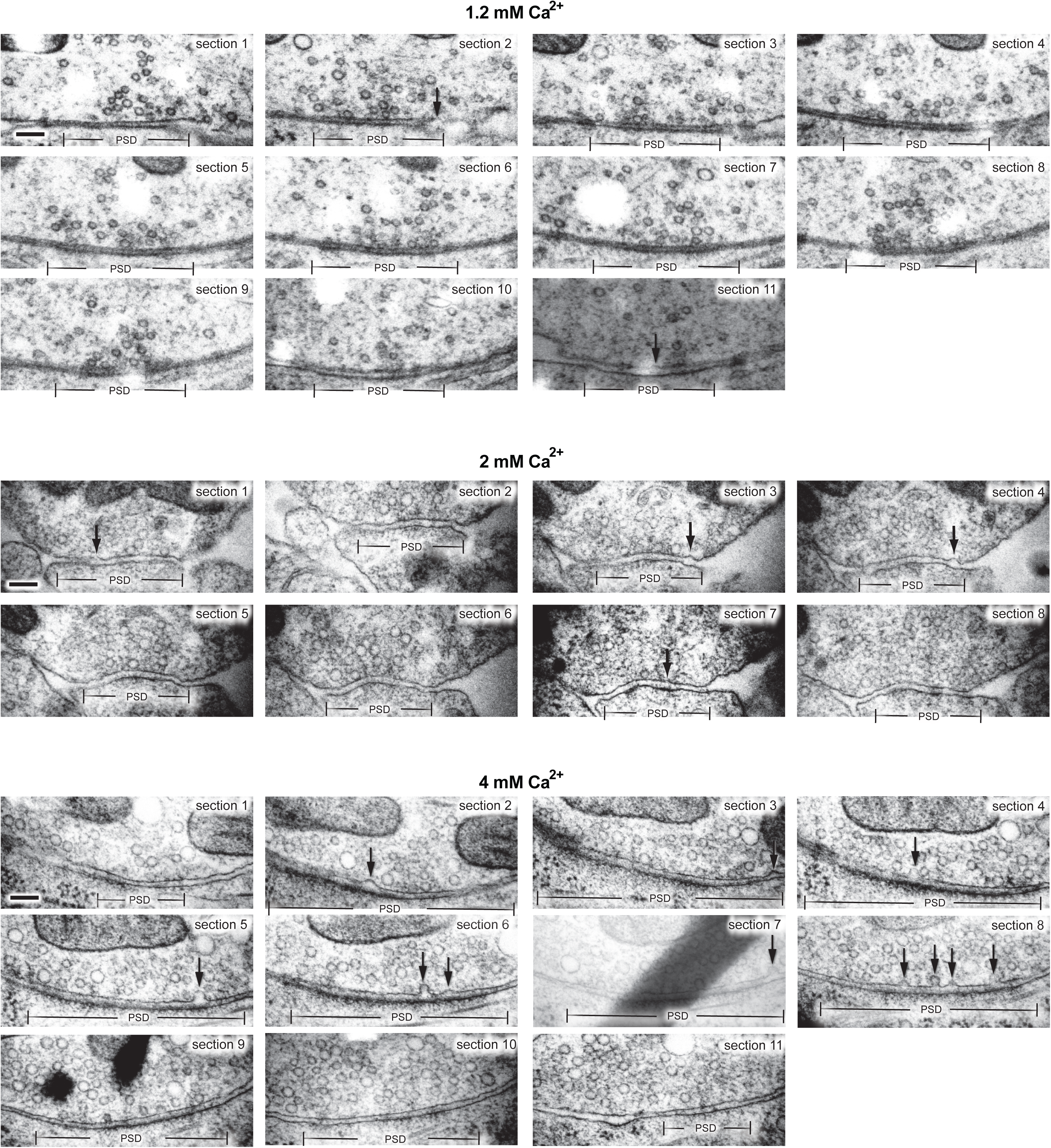
**Multiple fusion events in serial-sectioning reconstructions of active zones from neurons frozen 5 ms after an action potential. a**, Example transmission electron micrographs from serial sections of active zones from neurons frozen 5 ms after an action potential in 1.2 mM, 2 mM, and 4 mM extracellular Ca^2+^ (from the same experiments described in Figure 2). Scale bar: 100 nm. PSD: post-synaptic density. AP: action potential. Arrows indicate “pits” in the active zone (opposite the post-synaptic density), which are presumed to be vesicles fusing with the plasma membrane. Note that pits within the same active zone are often different depths.

**Supplementary Figure 3.**
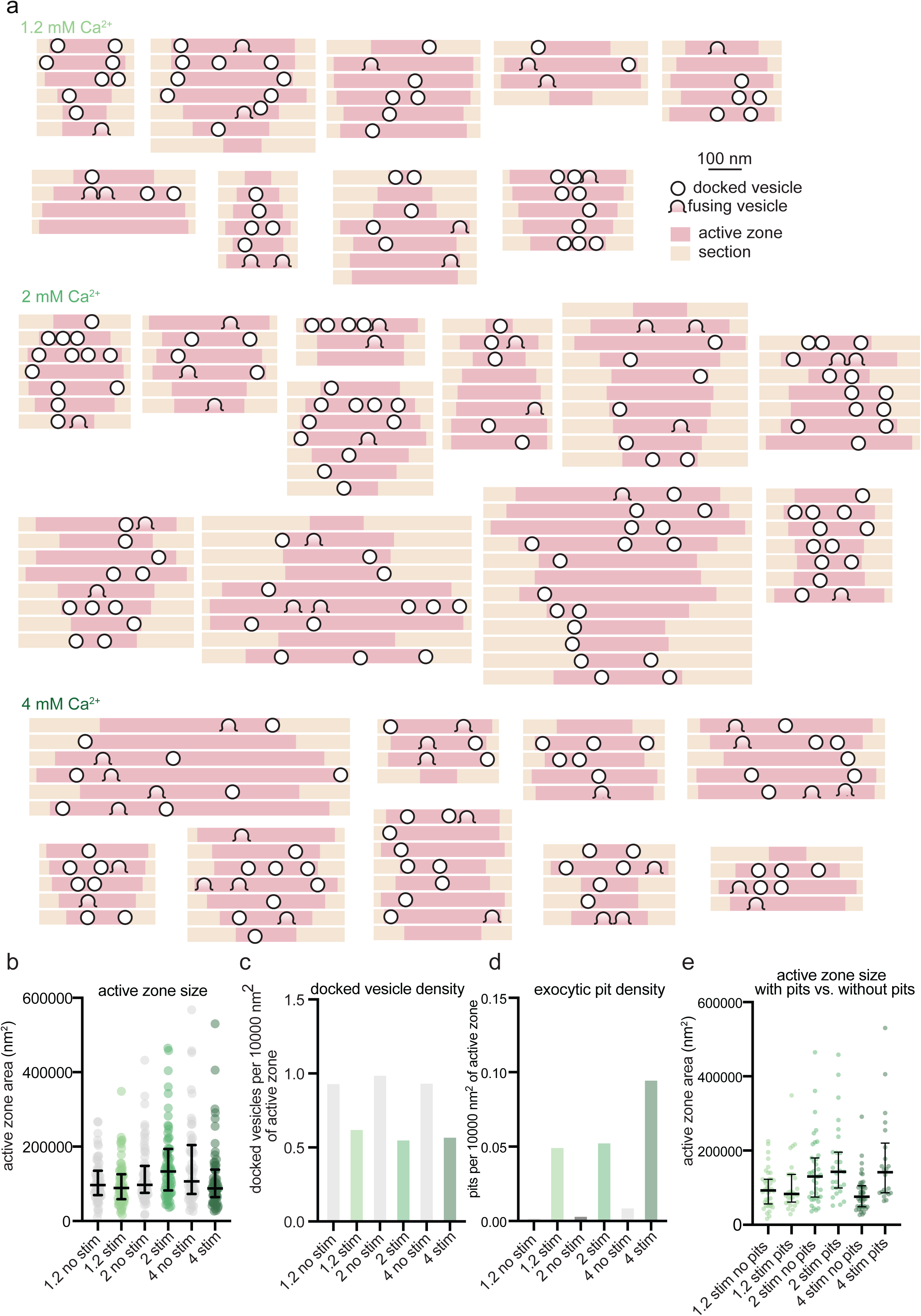
**Representations of serial-sectioning reconstructions of active zones with exocytic pits. a**, Graphical depictions of serial-sectioned active zones containing exocytic pits, from the experiments described in Figure 2. **b**, Sizes of active zones from the data set shown in Figure 2. Area was calculated as the product of the longest 2-D length of active zone in a 2-D profile from that synapse, the number of sections containing the active zone, and section thickness (that is, as the area of a rectangle). Error bars: median and interquartile range. **c**, Density of docked vesicles in the active zone, from the experiments shown in Figure 2. Calculated as the total number of docked vesicles from a given sample x10000, divided by the sum of the total area of active zones from that sample. **d**, Density of exocytic pits in the active zone, from the experiments shown in Figure 2. Calculated as the total number of pits from a given sample x10000, divided by the sum of the total area of active zones from that sample. **e**, Same data shown in **b**, sorted by whether the active zone contained an exocytic pit or not. See Supplementary Data Table 1 for full pairwise comparisons and summary statistics.

**Supplementary Figure 4.**
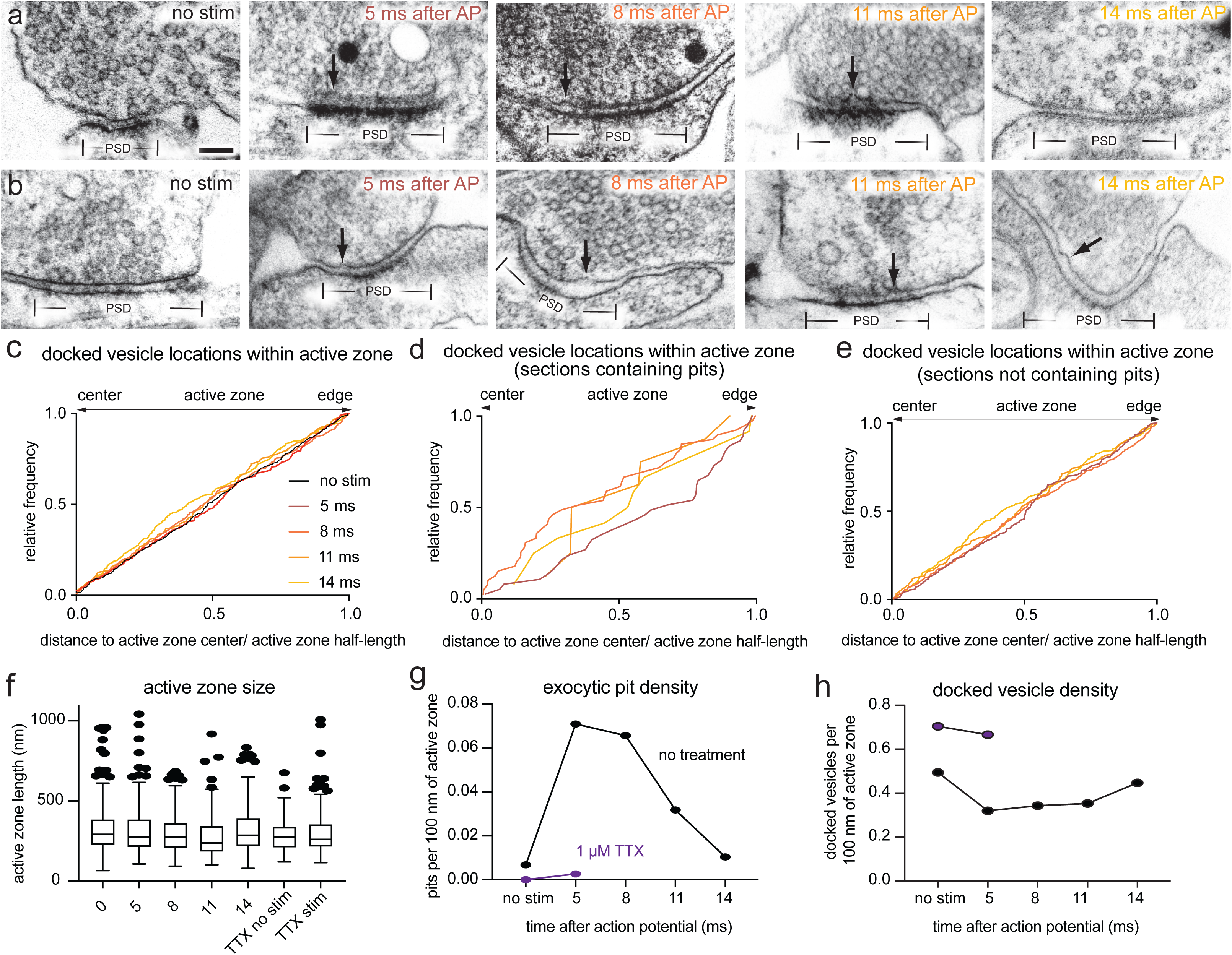
**Fusion intermediates at multiple time points during the first 14 ms after an action potential. a-b**, Example transmission electron micrographs of synapses from neurons frozen without stimulation or 5, 8, 11, or 14 ms after an action potential (these are other examples from the same experiments shown in Figure 3). **c**, Cumulative relative frequency of locations of docked vesicles within the active zone, normalized to the size of the active zone (no stim, n = 447; 5 ms, n = 300; 8 ms, n = 348; 11 ms, n = 188; 14 ms, n = 306 docked vesicles). **d**, Same data as in **c**, showing only vesicles from synaptic profiles that contain pits. **e**, Same data as in **c**, showing only vesicles from synaptic profiles that contain do not contain pits. **f**, Size of active zones from data in Figure 3. Box, median and interquartile range; whiskers, 25^th^ minus 1.5x interquartile range and 75^th^ plus 1.5x interquartile range; dots indicate values outside this range. **g**, Density of pits in the active zone, from the experiments shown in Figure 3. Calculated as the total number of pits from a given sample x100, divided by the sum of the total length of active zones from that sample. **h**, Density of docked vesicles in the active zone, from the experiments shown in Figure 3. Calculated as the total number of docked vesicles from a given sample x100, divided by the sum of the total length of active zones from that sample. Scale bar: 100 nm. PSD: post-synaptic density. AP: action potential. Arrows indicate “pits” in the active zone (opposite the post-synaptic density), which are presumed to be vesicles fusing with the plasma membrane. See Supplementary Data Table 1 for full pairwise comparisons and summary statistics.

**Supplementary Figure 5.**
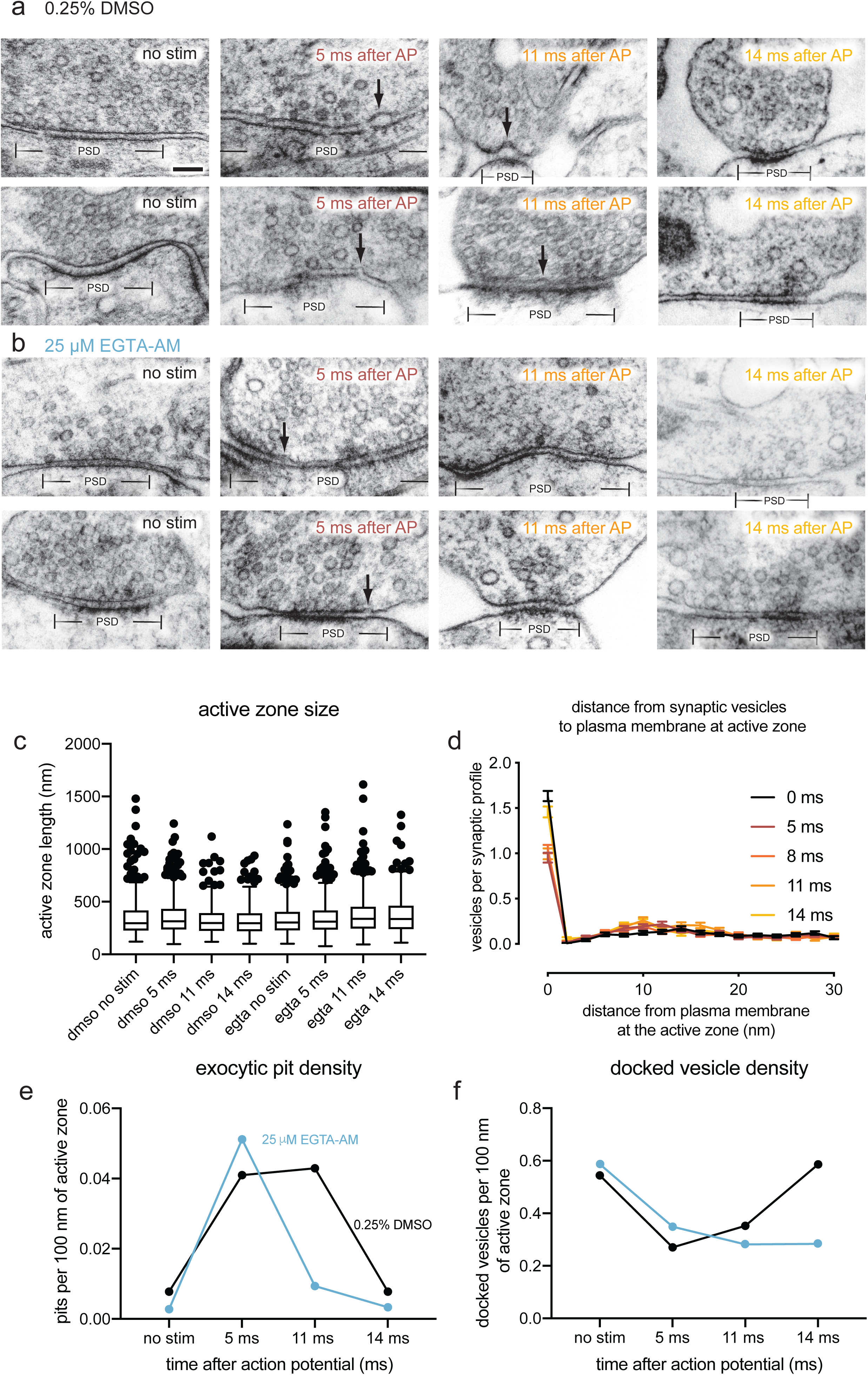
**Chelating residual Ca^2+^ blocks fusion intermediates at 11 ms but not 5 ms after an action potential. a-b**, Example transmission electron micrographs of synapses from neurons pre-treated with **a** 0.25% DMSO or **b** 25 μM EGTA-AM and frozen either without stimulation, 5 ms after stimulation, or 11 ms after stimulation (these are other examples from the same experiments shown in Figure 4). Scale bar: 100 nm. PSD: post-synaptic density. AP: action potential. Arrows indicate “pits” in the active zone (opposite the post-synaptic density), which are presumed to be vesicles fusing with the plasma membrane. **c**, Size of active zones from data in Figure 4. Box, median and 25-75^th^ percentile; whiskers, 25^th^ minus 1.5x interquartile range and 75^th^ plus 1.5x interquartile range; dots indicate values outside this range. See Supplementary Data Table 1 for full pairwise comparisons and summary statistics. **d**, Distances of synaptic vesicles from the plasma membrane at the active zone, including both vesicles that were annotated as docked and not docked. Distances are binned in 2-nm increments, except for “0”, which indicates vesicles ∼0 nm from the active zone membrane (“2” indicates vesicles 0.1-2 nm from the membrane, “4” indicates 3-4 nm, etc.).

